# Primed smooth muscle cells acting as first responder cells in disease

**DOI:** 10.1101/2020.10.19.345769

**Authors:** Matt D Worssam, Jordi Lambert, Sebnem Oc, Annabel L Taylor, Lina Dobnikar, Joel Chappell, Jennifer L Harman, Nichola L Figg, Alison Finigan, Kirsty Foote, Anna K Uryga, Martin R Bennett, Mikhail Spivakov, Helle F Jørgensen

## Abstract

**Rationale:** Vascular smooth muscle cell (VSMC) dysregulation is a hallmark of vascular disease, including atherosclerosis. In particular, the majority of cells within atherosclerotic lesions are generated from pre-existing VSMCs and a clonal nature has been documented for VSMC-derived cells in multiple disease models. However, the mechanisms underlying the generation of oligoclonal lesions and the phenotype of proliferating VSMCs are unknown.

**Objective:** To understand the cellular mechanisms underlying clonal VSMC expansion in disease.

**Methods and Results:** Here we analyse clonal dynamics in multi-color lineage-traced animals over time after vessel injury. We demonstrate that VSMC proliferation is initiated in a small fraction of VSMCs that initially expand clonally in the medial layer and then migrate to form the oligoclonal neointima. Selective activation of VSMC proliferation also occurs *in vitro*, suggesting that this is a cell-autonomous feature. Mapping of VSMC trajectories using single-cell RNA-sequencing reveals a continuum of cellular states after injury and suggests that VSMC proliferation initiates in cells that have downregulated the contractile phenotype and show evidence of pronounced phenotypic switching. We show that proliferation is associated with induced expression of stem cell antigen 1 (SCA1) and the expression signature previously identified in SCA1+ VSMCs in healthy arteries. A remarkably increased proliferation of SCA1+ VSMCs, directly validated in functional assays, indicates that SCA1+ VSMCs act as “first responders” in vascular injury. Early atherosclerotic lesions also had clonal VSMC contribution and we show that the proliferation-associated injury response is conserved in plaque VSMCs, extending these findings to atherosclerosis. Finally, we identify VSMCs in healthy human arteries that correspond to the SCA1+ state in mouse VSMCs and show that genes identified as differentially expressed in this human VSMC subpopulation are enriched for genes showing genetic association with cardiovascular disease.

**Conclusions:** We show that cell-intrinsic, selective VSMC activation drives clonal proliferation after injury and in atherosclerosis. Our study suggests that healthy mouse and human arteries contain VSMCs characterised by expression of disease-associated genes that are predisposed for proliferation. Targeting such “first responder” cells in patients undergoing vascular surgery could effectively prevent injury-associated VSMC activation and neoatherosclerosis.

## INTRODUCTION

Cardiovascular disease remains the leading cause of death worldwide according the World Health Organization. Current treatment strategies focus on lipid lowering and blood pressure regulation. However, recent genetic, cellular and molecular studies highlight an important role for vessel wall cells in disease development^1^. Vascular smooth muscle cell (VSMC) contractility controls vessel tone, but the cells display a remarkable phenotypic plasticity characterized by downregulation of the contractile machinery (e.g. *MYH11, ACTA2* and *CNN1*), concomitant increased expression of extracellular matrix (ECM) components (e.g. Matrix gla protein, MGP, and collagen) and exit from quiescence^2^. Such classical VSMC phenotypic switching ensures tissue homeostasis and enables physiological vessel remodeling, but deregulated VSMC plasticity in cardiovascular disease contributes to lesion development and arterial remodeling after surgical intervention^1,3,4^.

VSMC-derived cells constitute the major cellular component of atherosclerotic lesions^1,3-5^. Multi-color lineage tracing studies in mouse models show that lesional VSMCs are oligo-clonal and generated from very few pre-existing VSMCs^6-10^. Similar clonal VSMC contribution has also been observed in aortic dissection^11^ and after injury^7^. The oligoclonal nature of intimal cells appears at odds with observations suggesting widespread VSMC proliferation after injury^12-14^, calling for analysis of whether lesional clonality of VSMC-derived cells results from selective activation of a small number of VSMCs or if this is due to differential survival of VSMC clones following a general proliferative response (discussed in Liu&Gomez^9^).

Using single cell-RNA sequencing (scRNA-seq), we and others have delineated VSMC heterogeneity in atherosclerotic lesions, including the demonstration of substantial variability in expression of genes associated with cardiovascular disease^14-19^. However, the diversity of VSMCs in lesions does not inform about how VSMC investment in plaques is initiated, particularly with regard to the resulting clonal outcome. We previously identified a small subset of VSMCs in healthy mouse vessels marked by stem cell antigen 1 (SCA1), that have reduced contractile gene expression and show increased expression of genes associated with cell activation^15^. SCA1 is also induced in VSMCs in atherosclerosis and other disease models^15-17^, suggesting that this VSMC subset is relevant for disease development. Whether these atypical cells are functionally distinct is not known and the presence an equivalent human cell population in healthy arteries remains to be demonstrated.

By studying clonal dynamics and acute changes in scRNA-seq profiles in mouse models we here provide evidence that cell-intrinsic mechanisms, associated with a primed cellular state characterized by SCA1 expression, result in selective proliferation of a small number of VSMCs in both atherosclerosis and after injury. We further demonstrate an 8-fold increased proliferative capacity of SCA1-expressing VSMCs from healthy animals, suggesting that these act as “first responder” cells that will rapidly start proliferating after insults. An equivalent VSMC subpopulation displaying the primed gene expression signature is identified through scRNA-seq analysis of healthy human arteries. Collectively, our data suggests that selective activation of predisposed VSMCs could underlie the development of human atherosclerosis and that targeting these cells could represent novel therapeutics in atherosclerosis prevention.

## METHODS

(Extended methods are available in the Online-only Data Supplement).

### Human tissue

Anonymized plaque-containing carotids and normal human aorta tissue was obtained from patients undergoing carotid endarterectomy (50F, 75M, 74M, 54M) or either coronary artery bypass or valve replacement (52M, 63F, 42M, 60M, 85M, 74M) respectively, under informed consent using protocols approved by the Cambridge or Huntingdon Research Ethical Committee.

### Animals and procedures

Animal experiments were approved by the local ethics committee and performed according to UK Home Office regulation under project license P452C4595. All alleles have been described previously; Myh11-CreERt2 (Myh11) confers expression of a tamoxifen-inducible Cre recombinase in smooth muscle cells^18,19^, Rosa26-Confetti (Confetti)^20^ and Rosa26-EYFP (EYFP)^21^ are Cre-recombination reporter alleles, Ki67/RFP is an insertion in the Mki67 locus resulting in expression of a KI67/RFP fusion protein^22^ and the mutant Apoe allele sensitizes mice to high fat diet (HFD)-induced atherosclerosis development^23^. VSMC lineage labeling was achieved by intraperitoneal tamoxifen injections (10x 0.1 mg tamoxifen over 2 weeks) followed by at least 1 week rest period for tamoxifen metabolism. Only male animals were used as the Myh11-CreERt2 transgene is Y-linked. The left carotid artery was ligated under the bifurcation with a silk suture under anesthesia (2.5-3% isofluorane by inhalation) with subcutaneous pre-operative analgesic (∼0.1 mg/kg body weight, Buprenorphine) as described^7^. HFD (Special Diets Services, containing 21% fat and 0.2% cholesterol) was administered for 9-24 weeks.

### Tissue analysis

Arteries were fixed, processed for whole mount and confocal imaging followed by cryo-sectioning, or directly cryo-sectioned and stained as described^7^. Aortic tissue explants were injured using a forceps pinch and embedded in Matrigel. Single cell suspensions were generated by enzymatic digestion (Collagenase IV, Invitrogen and Elastase, Worthington) for scRNA-seq library preparation, ImageStream analysis, flow cytometry and *in vitro* culture. Antibodies used for staining are described in Table I of the Online-only Data Supplement.

### Single cell-RNA sequencing

Mouse scRNA-seq datasets were generated from VSMC-lineage labeled cells isolated by flow-assisted cell sorting (FACS) from ligated left carotid arteries of Myh11-EYFP-Ki67/RFP animals 5 (D5) or 7 days (D7) after surgery, using the 10x chromium system (mouse D5, mouse D7). The Smart-seq2 protocol was used to process index-sorted cells from Myh11-EYFP-Ki67/RFP animals 7 days after surgery and control unligated animals. Human scRNA-seq data were generated from medial cells isolated from a healthy aorta (65-year-old male). Datasets were analyzed using the CRAN R package Seurat (v.3.1.2)^24,25^ in *R* (v.3.6.2). Differential expression analysis was done using DEseq2^26^. Trajectory-inference was done using the *R* package slingshot (v.1.4.0)^27^. Summarized expression level of gene subsets was calculated and displayed as described^15^. Scripts used for data analysis are available upon request.

### Data accessibility

The scRNA-seq datasets generated in this study (mouse D5 10x, mouse D7 10x, mouse D7 Smart-seq2, human 10x) have been deposited in the Gene Expression Omnibus (GEO) repository; accession number will be made available prior to publication. The scRNA-seq dataset of VSMC-lineage-labeled plaque cells from high fat diet-fed Myh11-Confetti-Apoe animals is available from GEO (accession number: GSE117963).

### Statistical analysis

Statistical analysis was performed in R, the Shapiro–Wilk test was used to ascertain normal distribution, equal variance assessed using Bartlett or Levine tests. Tests used to assess statistical significance are indicated in figure legends. Local regression analysis was used to fit a LOESS curve of patch number. To assess statistical significance of SCA1 expression status in the clonal proliferation assay, a generalized linear model was fitted for patch number whereas multiple linear regression used for patch area, as the data showed equal variance and linearity and the residuals were approximately normally distributed.

## RESULTS

### Proliferation is restricted to a small subset of VSMCs after injury

To determine whether the observed oligoclonal VSMC contribution to vascular disease^6,7,11^ results from selective cell activation or clonal competition following general activation of VSMC proliferation^9^ we employed the carotid ligation injury model. We previously showed that 28 days after surgery, neointimal lesions are composed of large clones derived from pre-existing VSMCs^7^, and the acute, reproducible induction of VSMC proliferation^5,14^ makes this model ideal for assessing the dynamics of such VSMC clone generation. Myh11-CreERt2/Rosa26-Confetti+ (Myh11-Confetti) animals were used to lineage-label VSMCs with one of 4 fluorescent proteins in a random fashion prior to surgery. We assessed the VSMC injury-response at different timepoints after ligation by whole-mount confocal microscopy (Figure 1A). Neointimal lesions were observed in most (15/16) animals analyzed from 12 days after surgery (Figure 1B, Table II and Figure I in the Online-only Data Supplement), consistent with previous findings^14^, and similar to day 28, these had oligoclonal VSMC contribution. Contiguous patches of lineage-labeled VSMCs of a single color were also identified in the medial layer of animals in 36 out of 43 injured arteries (Figure 1C), whereas none were detected in control animals. No patches were observed prior to day 5 (D5), where 2 of 3 arteries contained small medial patches (Figure 1C, D and Table II in the Online-only Data Supplement). Formation of medial VSMC patches prior to neointimal invasion is in keeping with previous observations that medial proliferation precedes neointimal formation^14^. Quantification of patches and their size distribution per artery demonstrated a gradual increase in both number and size over time, with a possible plateau after 2 weeks (Figure 1B-D). Interestingly, the number of intimal VSMC clones was much lower than medial patch numbers, suggesting that neointimal lesions are generated by a subset of the VSMCs that activate proliferation (Figure 1B, Figure IC in the Online-only Data Supplement). This is consistent with a model where only a few VSMCs initiate proliferation, whereas no evidence of clonal competition was observed.

**Figure 1:**
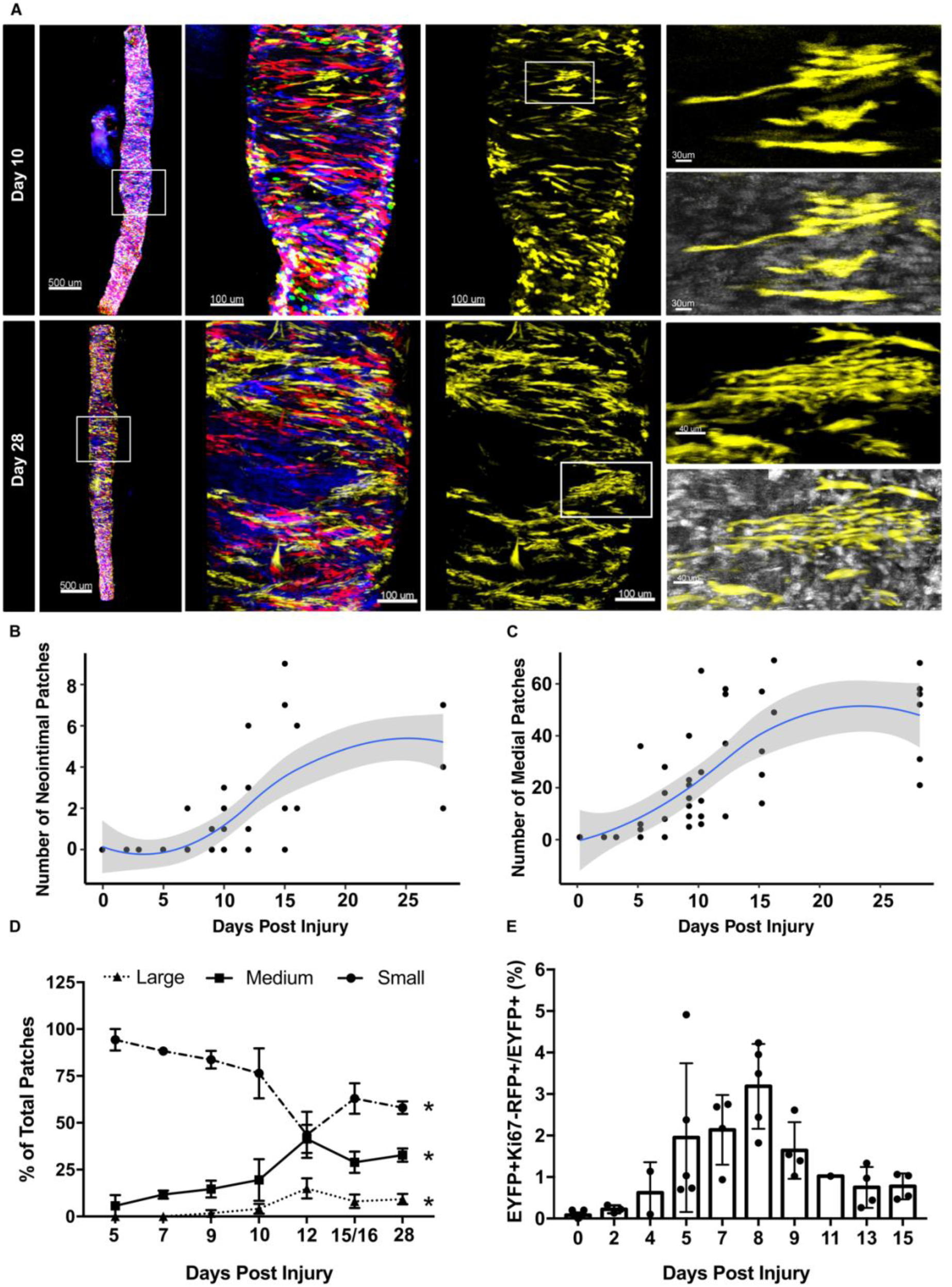
Analysis of clonal dynamics after vascular injury. **A**, Representative examples of whole-mounted left carotid arteries after ligation surgery in VSMC lineage-labeled Myh11-Confetti animals. Left panels show signals for all Confetti proteins in the whole vessel (scalebar = 500 μm) and a magnified segment of arteries (scalebar = 100 μm) analyzed at day 10 (top) and 28 (lower) after surgery. Right panels only show signals for the Confetti yellow fluorescent protein (YFP)-reporter for the magnified arterial segment. Far-right panels show YFP only (top) and YFP+DAPI (lower) in magnified views of boxed regions, with VSMC singlets and small (top, Day 10, scalebar = 30 μm) and large VSMC patch (lower, Day 28, scalebar = 40 μm). **B-D**, Quantification of neointimal (B) or medial VSMC patch numbers (C) and the size distribution of medial patches (D) over time after injury in a total of 43 arteries. **B, C**, Dots show values for individual arteries, blue line shows local polynomial regression (LOESS) regression and grey area represents the 95% confidence interval. **D**, The fraction of small (circles), medium (squares) and large patches (triangles) are shown for each analysis time point. Mean and S.E.M. for each timepoint are indicated. Asterisks indicate statistically significant change over time for all size groups (P<0.05) according to a Kruskal-Wallis test. **E**, The fraction of EYFP+ cells expressing the Ki67/RFP reporter in a FACS analysis of VSMCs from Myh11-EYFP-Ki67/RFP animals. Dots show values for individual arteries (32 total), bars represent mean and error bars indicate standard deviation.

Further supporting the idea that activation of proliferation is limited to a subset of VSMCs, EdU incorporation was rarely detected in lineage-labeled VSMCs in D5 arteries (Figure ID in the Online-only Data Supplement). Low VSMC proliferation frequency was confirmed by FACS analysis of Myh11-CreERt2;Rosa26-EYFP;Ki67/RFP (Myh11-EYFP-Ki67/RFP) animals, where tamoxifen injection results in EYFP+ VSMCs and where proliferating cells express a Ki67-RFP reporter protein^22^ (Figure 1E). Double positive (EYFP+RFP+) cells were almost absent in healthy arteries, increased in frequency from D5, peaked at D8 and never constituted more than 5% of all EYFP+ VSMCs.

Collectively, these data suggest that activation of VSMCs proliferation occurs at a low frequency and that only a fraction of medial VSMC clones migrate across the inner elastic lamina to form oligoclonal lesions.

### VSMC investment in atherosclerotic plaques mimics the injury response

Almost all medial and intimal VSMC patches were restricted to arterial segments with increased medial diameter. These remodeled segments or “bulges”, also displayed disorganized cellular arrangement (Figure 2A-C, Figure IE in the Online-only Data Supplement). Such remodeled segments were observed in all arteries displaying patches, as well as in 2/5 arteries prior to day 5 (Table I in the Online-only Data Supplement) and did not increase in length over time (Figure 2B). Despite the onset of VSMC proliferation, reduced medial cell density was observed at early timepoints in injured arteries compared to healthy controls (Figure 2C), in accordance with previous observations^14,28^. Evidence of low cellularity, defined as absence of both Confetti signal and nuclear staining, was more pronounced in bulged regions compared to arterial segments without remodeling (Figure 2C, D). This suggests that VSMC death and matrix remodeling occurs at the same time, or precedes, VSMC proliferation. Notably, at all timepoints, a large number of “singlet” VSMCs persisted, even within the bulged regions (Figure 2A, Figure IA, B in the Online-only Data Supplement), demonstrating that despite a shared signaling environment, not all VSMCs within bulges exit quiescence and initiate clonal expansion.

**Figure 2:**
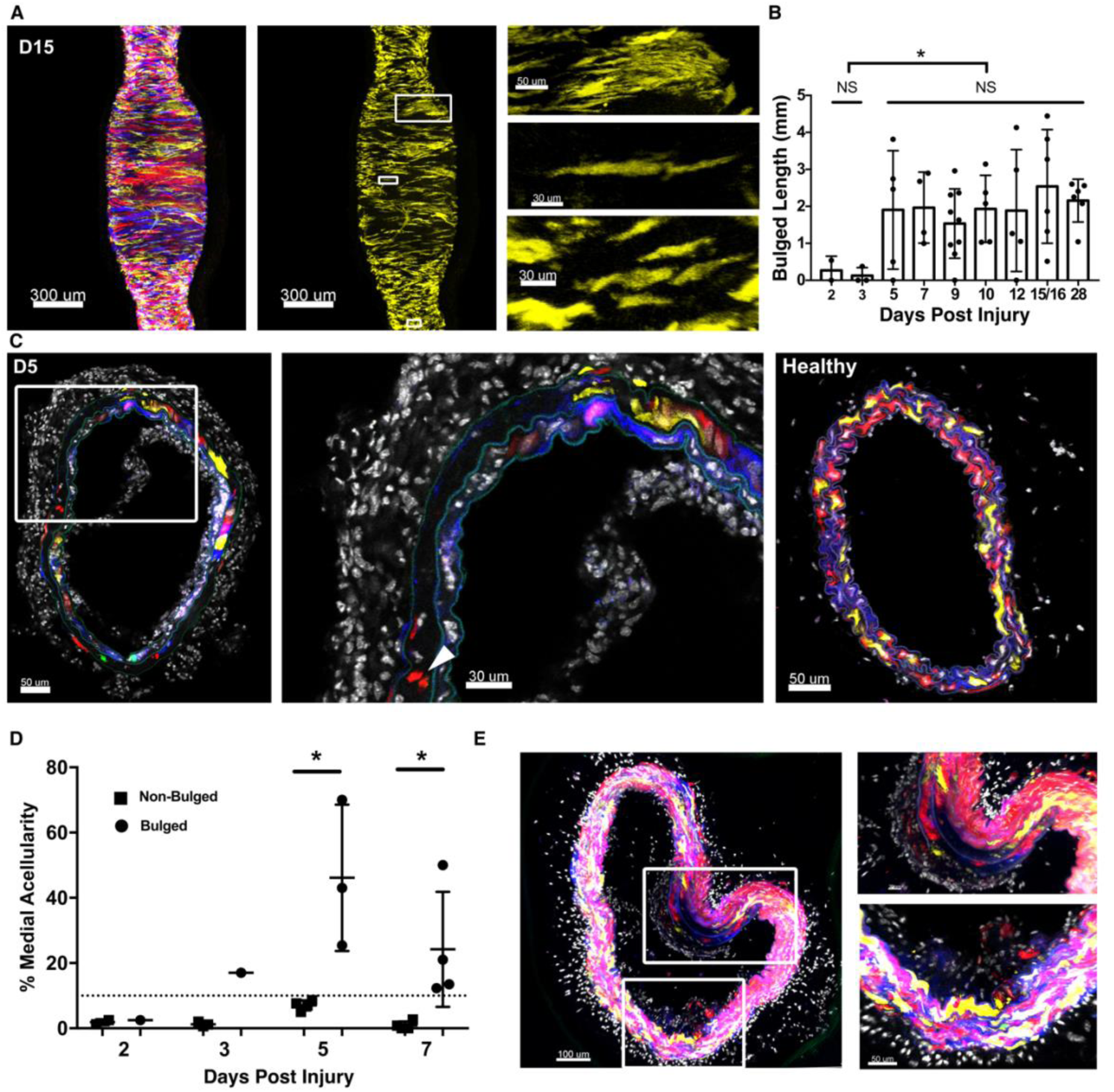
Early atherosclerotic plaques share the selective VSMC proliferation and remodeling of the vessel wall observed after injury. **A**, Confocal image (max projection) of Myh11-Confetti carotid artery analyzed 15 days (D15) after ligation, showing co-localization of patches with “bulged” regions of increased medial diameter. Signals for all Confetti proteins (left) or yellow fluorescent (YFP) only (middle) are shown (scalebar = 300 μm). Right panels show magnified views of boxed regions with examples of a patch (top) or a non-patch VSMC singlet (middle) in a bulged region, and several VSMC singlets in an un-remodeled, non-bulged arterial segment (lower right panel). Scalebar = 30 μm. **B**, Length of bulged region in carotid arteries analyzed at indicated times after ligation. Dots show bulge length in individual arteries (43 total), bars show mean and error bars indicate standard deviation. Asterisks indicate P<0.05, Welch’s t-test. **C**, Cross-section of carotid arteries analyzed 5 days after ligation (left) or from unligated animal (right). Scalebar = 50 μm. Middle panel shows magnified view of boxed region from D5 artery (scalebar = 30 μm). Arrows point to medial VSMC patches. Signals for Confetti and DAPI are shown. **D**, Percentage of medial area without Confetti and DAPI signal in virtual cross-sections from whole mounted arteries 2-7 days after ligations (see Figure IE in the Online-only Data Supplement). Cross-sections from bulged (circles) or non-bulged regions (squares) were analyzed separately (each dot shows mean of 3-5 cross-sections per artery). Asterisks indicate P<0.05, Mann-Whitney U test. **E**, Representative confocal image (single Z-plane), showing VSMC contribution to early lesions in lineage-labeled Myh11-Confetti-Apoe animals (n=5, 9-15 weeks high fat diet). Confetti and DAPI signals are shown. Magnified views of boxed regions are shown on the right. Scalebar = 100 μm (left), 30 μm (top right) or 50 μm (lower right).

To assess whether selective VSMC proliferation also underlies oligoclonal VSMC contribution observed in atherosclerosis^6,7^, we analyzed Myh11-Confetti animals on an Apoe^-/-^ background (Myh11-Confetti-Apoe) with tamoxifen-mediated VSMC-lineage-labeling prior to feeding an atherogenic diet for 9-15 weeks. Analysis of early stage plaque (<50 Confetti+ cells per section) revealed VSMC investment to lesions from carotid arteries, aortic root, arch and the descending aorta (Figure 2E). Like late stage lesions^7^, VSMC-derived cells were typically only of a single color and where several Confetti colors were detected, cells were arranged in a non-random manner (Figure 2E). Examples of individual VSMC clones contributing exclusively to either cap or core was observed, and VSMC investment was found in lesions that lacked an obvious fibrous cap structure (Figure 2E). The medial layer generally remained mosaic with respect to Confetti protein expression, however, evidence of medial VSMC patches expressing the Confetti color observed in lesion VSMC clones was often observed underneath the plaque, where medial cell disarray and breaks in the elastic lamina were also detected (Figure 2E). Collectively, this analysis suggests that selective activation of VSMC proliferation and other hallmarks of VSMC injury-responses are also found in early atherogenesis.

### Selective initiation of VSMC proliferation *in vitro*

To investigate whether the selective clonal expansion observed *in vivo* is intrinsic to VSMCs, we cultured aortic tissue explants *in vitro* after introducing a forceps-pinch “injury” to promote VSMC proliferation (Figure 3A). After culture, tissue explants displayed persistent mosaic labeling in most regions but developed monochromatic patches similar to those observed *in vivo* along tissue edges and at forceps pinch-injuries (Figure 3B). Quantification of contiguous surfaces expressing the same Confetti protein (Figure II in the Online-only Data Supplement) confirmed that the vast majority of VSMC surfaces remained comparable in size to those detected in freshly isolated vessels (Figure 3C). On average, 30.5 (±10.8, S.E.M.) large VSMC patches were observed in 1 mm^2^ tissue explants after 8 days of culture, whereas these were rare at day 0 (2.2±1.3, Figure 3D). Confirming the idea that VSMC patches resulted from selective proliferation of a small number of cells, similarly sized patches were observed in tissue explants from animals with reduced labeling density (Figure II in the Online-only Data Supplement).

**Figure 3:**
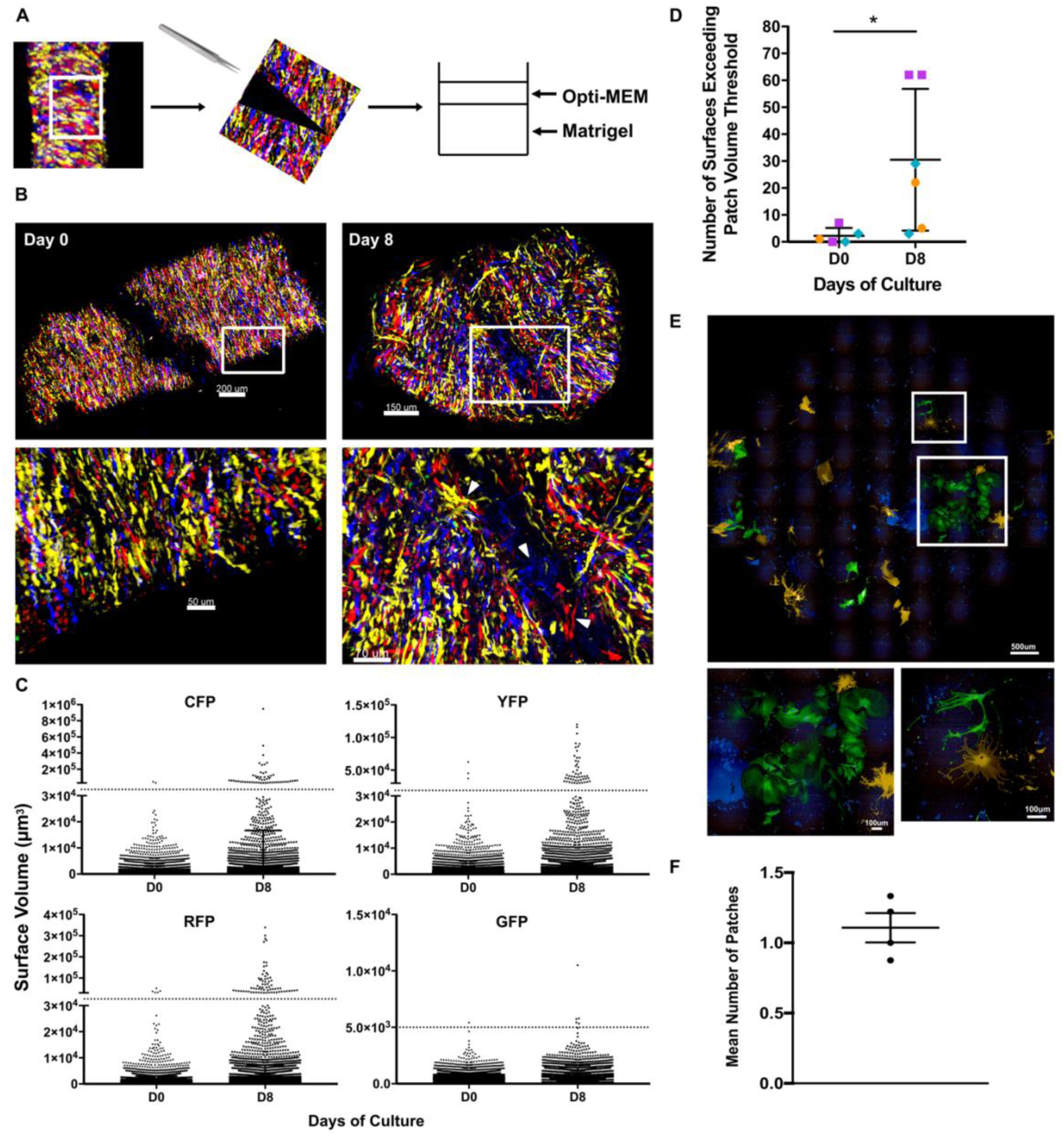
Selective activation of VSMC proliferation *in vitro*. **A**, Schematic of tissue explant forceps-pinch-injury-assay. **B**, Confocal images (max projection) of tissue explants before (Day 0, left) or after (Day 8, right) culture. Magnified views of boxed regions are shown in lower panels, where arrowheads point to monochromatic VSMC patches within or adjacent to forceps pinch injury. Scalebar = 200 μm (top left), 150 μm (top right), 50 μm (lower left), 70 μm (lower right). **C-D**, Quantification of Confetti+ regions in tissue explants before (D0) or after (D8) culture, done using surface-rendering in Imaris. **C**, volumes of individual surfaces are shown separately for each Confetti protein. Thresholds for day 0 surface volumes indicated by dotted lines. **D**, the number of surfaces exceeding day 0 thresholds is shown for each explant. Symbols and colors indicate explants from the same animal. Three animals (1-2 explants per timepoint from each animal) were analysed. Asterisks indicate P<0.05, Welch’s t-test. **E**, Confocal live cell image of VSMCs from Myh11-Confetti and wild type animals (mixed 1:3) after 2 weeks of culture (top). Lower panels show magnified views of boxed regions with representative VSMC patch (lower left) or non-expanded singlet Confetti+ cells (lower right). **F**, Quantification of number of Confetti+ patches per well. Dots show mean of 3 wells per animal. Mean and S.E.M. from 4 animals, analyzed separately, are shown. Scalebar = 500 μm (top), 100 μm (lower panels).

To test whether selective proliferation also occurred in freshly isolated, enzymatically dissociated VSMCs, we designed an assay that allowed detection of emerging VSMC clones while maintaining the cell-cell contacts required for VSMC survival. Single cell suspensions of lineage-labeled VSMCs from Myh11-Confetti animals were mixed with wild-type VSMCs and live cell imaging performed periodically over a 3-week period (Figure 3E). Most Confetti+ cells in these cultures remained as “singlets” isolated by wild-type cells but occasionally a small patch of lineage-labeled VSMCs of one color formed (Figure 3F); 1.1 (±0.1 S.E.M) patch-forming cells out of 1250 Confetti+ VSMCs seeded.

Collectively, the selective proliferation of a small fraction of VSMC-lineage labeled cells in vitro indicates that activation of VSMC proliferation is a rare, cell-autonomous event.

### A continuum of VSMC expression profiles are observed after injury

To assess whether the observed selective VSMC proliferation reflects a heterogeneous injury response suggestive of distinct VSMC subpopulations, we generated single cell-RNA-seq (scRNA-seq) profiles of VSMCs 7 days after injury. For this analysis, EYFP+ VSMCs were isolated from Myh11-EYFP-Ki67/RFP animals and enriched for proliferating, double positive (EYFP+RFP+) cells by FACS. After quality control to remove cells with high mitochondrial read percentage and low number of detected genes, the profiles of 1126 cells were clustered and assessed for expression of markers of VSMC phenotypic state^29^ and genes associated with VSMC-derived plaque cells (*Lum, Tnfrs11b*^17^)(Figure 4A, B). A small group of cells (23 cells; cluster 11) clustered separately from the bulk of the population, expressed *Myh11* and other contractile genes, and no-to-low levels of proliferation markers (Figure III in the Online-only Data Supplement). The remaining cells formed a single structure and selected marker genes showed overlapping expression domains with gradually varying levels across the population; high expression of contractile genes (*Myh11*) in cell clusters 1, 2, 3 and 6, “synthetic” genes (*Mgp, Spp1, Col8a1*) peaked in clusters 0, 5, 7, 9 and 4 and proliferation markers (*Mik67*) were restricted to cells in clusters 8 and 10 (Figure 4B). Interestingly, the expression domain of *Ly6a*, encoding SCA1 and expressed in rare cells prior to injury^15^, was immediately adjacent to *Mki67*+ cells in cluster 10. Similar overlapping gene expression gradients across the cells were observed for cluster markers (Figure 4C, Table III in the Online-only Data Supplement) confirming that rather than being formed by discrete cellular subsets, lineage-labeled cells represented a continuum of varying cell states. We merged this unselected, but proliferation-enriched, dataset with profiles of cells index-sorted for the VSMC-lineage label (EYFP) and the Ki67/RFP reporter from injured vessels, or EYFP+ cells from control animals (Figure 4D). This suggested that VSMCs post-injury display a spectrum of phenotypes from a contractile *Myh11*-positive state similar to that seen in healthy vessels (cell clusters 1, 2, 3 and 6), to a proliferative state characterized by high S and G2M scores (Figure 4A, B). In addition to increasing levels of classical markers of a synthetic VSMC state, the total number of genes detected also increased gradually along this axis (Figure 4B) indicative of increasing activation level. These analyses show that the transcriptional signatures of proliferating VSMCs substantially overlap with that of non-proliferating VSMCs after injury, suggesting that cells adopt states along a trajectory from a quiescent-contractile to a proliferative state.

**Figure 4:**
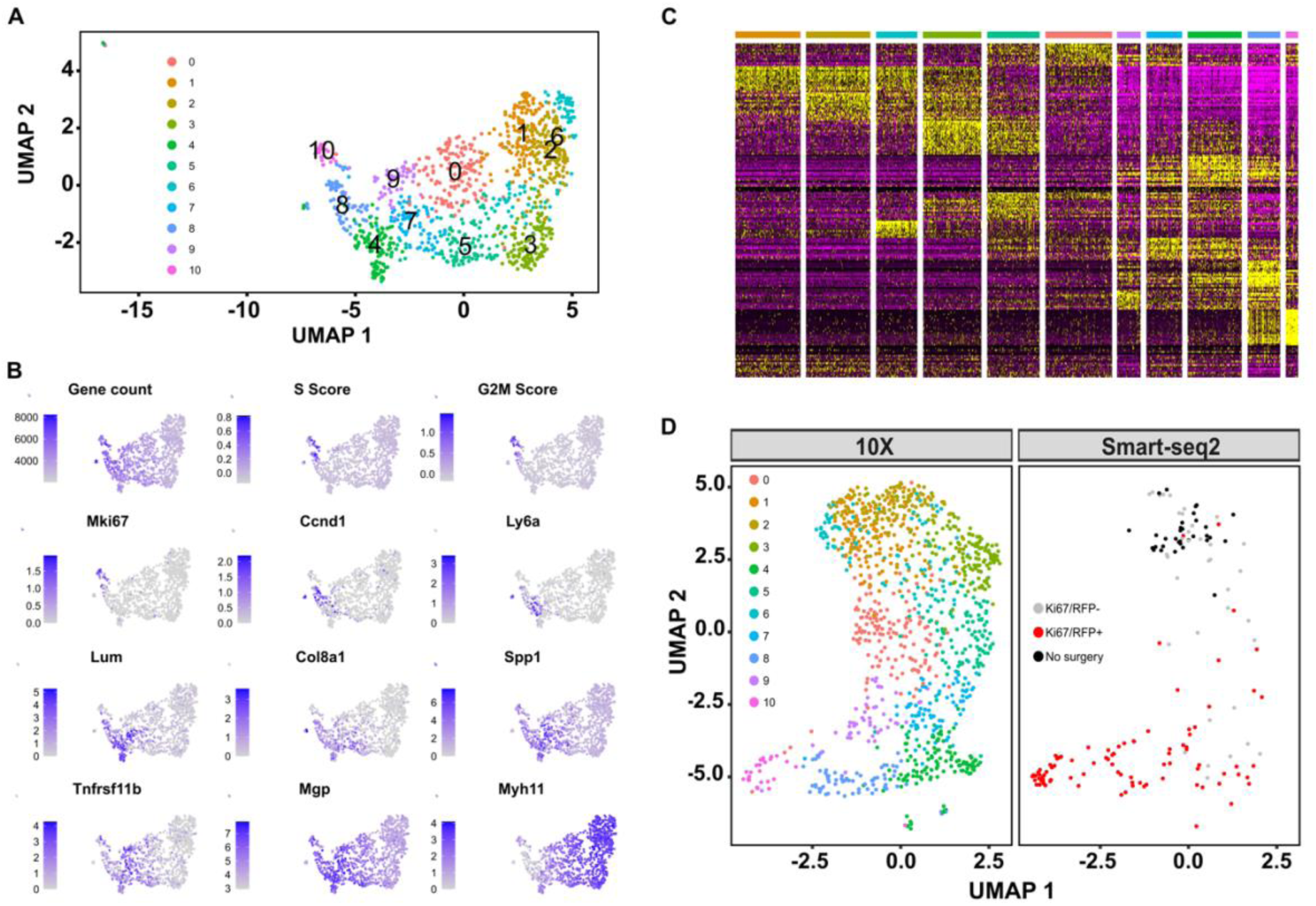
Gradual changes from the contractile to a proliferation-associated expression signature. **A**, Uniform Manifold Approximation and Projection (UMAP) showing the cell cluster map for scRNA-seq (10x Chromium) analysis of 1103 VSMC-lineage label positive cells from ligated left carotid arteries 7 days after surgery. A cell cluster representing a minor VSMC population (cluster 11, 23 cells) is not shown, see Figure III in the Online-only Data Supplement. **B**, UMAP plots showing gene count, cell cycle scores and expression level of marker genes using grey-blue scales. **C**, Heatmap showing expression of the top cluster markers using a scale from purple (low) to yellow (high). Cells are arranged according to clusters as identified in top bar using color scale from panel A. **D**, UMAP of integrated dataset (10x and Smart-seq2 datasets, both Day 7), split by dataset. Colors show cell cluster identity (10x dataset, left) and index-sort identity (Smart-seq2 dataset, right) with cells from unligated control animals in black, and cells from ligated left carotid arteries 7 days after surgery in red (Ki67/RFP+) or grey (Ki67/RFP-).

### Evidence of segregated injury responses at the onset of VSMC proliferation

Our analysis suggests that VSMC proliferation does not result from activation of a distinct subpopulation of VSMCs, but rather derives from cells displaying extensive phenotypic switching. To investigate this idea further, we profiled cells 5 days after injury at the onset of VSMC proliferation (Figure 1E). Similar to day 7, VSMCs formed a continuous population displaying anticorrelated gradual changes in contractile and synthetic markers, with *Mki67* expression restricted to cell cluster 9 (Figure 5A). Surprisingly, trajectory inference using the Slingshot algorithm^27^, suggested the existence of two distinct VSMC injury-responses, of which only one included proliferating cells in cluster 9 (Figure 5B), and partition-based graph abstraction (PAGA)^30^ analysis confirmed these trajectories (Figure IV in the Online-only Data Supplement). As shown in Figure 5B, the pseudotime axes defining these two paths shared a common origin in the *Myh11*-positive cell clusters. Genes showing significant changes in expression (p-adj<0.05, log(fold change)>0.5) along pseudotime for Path1 or Path2 were identified using generalized additive models (GAMs) and organized into gene clades based on Pearson correlation (Figure 5B, lower panels, and Table IV in the Online-only Data Supplement). Contractile genes (e.g. *Myh11* and *Acta2*) and actin cytoskeleton organizers (e.g. *Rock1* and *Lmod1*) reduced expression along both pseudotime axes (Figure 5B, C and Table IV in the Online-only Data Supplement). Genes with increased expression along the trajectories also showed substantial overlap as well as some significant differences (Figure 5C and Table IV in the Online-only Data Supplement). Genes associated with cell cycle regulation, such as *Ccnd1* and *Mki67* were specifically induced along Path1, whereas Path2-induced genes were enriched for gene ontology (GO)-terms associated with protein folding (Figure V and Table V in the Online-only Data Supplement). Several heat shock genes were upregulated along both Path1 and Path2 (Figure 5B, C). To verify these findings and rule out dissociation-associated gene-expression changes^31^, we immunostained day 5 cryosections for the heat-shock factor alpha-crystallin B chain (CRYAB). CRYAB was detected in ∼30% of Confetti+ cells in bulged regions (Figure 5D, n=3), whereas <7% of Confetti+ cells in non-bulged regions of injured vessels were CRYAB+, similar to levels observed in control, unligated animals (2%). This confirms the *Cryab* upregulation observed by scRNA-seq and demonstrates a regional injury-associated VSMC response in bulged regions. Many *bona fide* cell cycle genes, such as *Pcna* and *Top2a*, were restricted to the Mki67+ cell cluster 9 and mapped to Path1 gene clade 2 (Table IV in the Online-only Data Supplement). In contrast, the expression domains of other Path1-induced genes (gene clades 1, 5, 7, 8, 10) were broader and many overlapped both cells in cluster 9 and pre-proliferative cells in cluster 3 (e.g. *Fbln2, Lum, Ccnd1*, Figure 5C). GO-term analysis of Path1-induced genes revealed enrichment for genes associated with collagen biosynthesis and fibril formation (e.g. *Col1a2, Col5a1* and *Errfi1*), extracellular matrix organization (e.g. *Fbln2, Fn1, Lum*), response to wounding (e.g. *Fgf2, Pdgfa*) and substrate adhesion (e.g. *Cdh13, Vcam1*), in addition to cell-cycle associated genes (Figure V and Table V in the Online-only Data Supplement).

**Figure 5:**
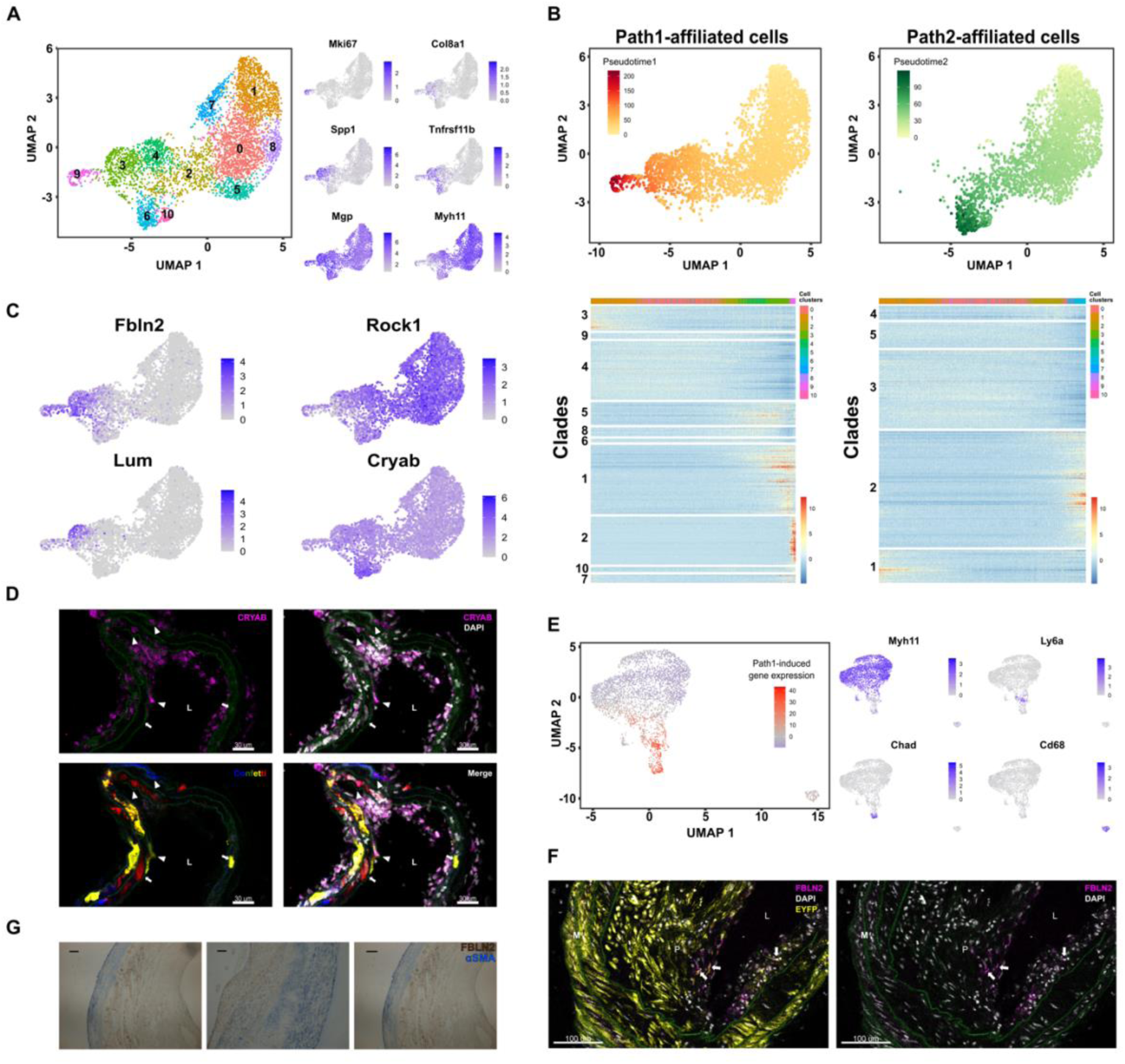
Identification of genes associated with onset of VSMC proliferation. **A**, Uniform Manifold Approximation and Projection (UMAP) showing cluster map (left) and expression level of marker genes using a grey-blue scale (right) in a scRNA-seq (10x Chromium) dataset of 4469 VSMC-lineage label positive cells from ligated left carotid arteries 5 days after surgery. **B**, Top panels are UMAPs showing only cells that are part of Path1 (left) or Path2 (right) with pseudotimes for each path indicated using a yellow-red (Path1, left) or yellow-green scale (Path2, right). Lower panels show heatmaps of genes with significant path-associated expression (p-adj<0.05, log(fold-change)>0.5), clustered into 10 (Path1, left) or 5 (Path2, right) gene clades. **C**, UMAP showing expression of trajectory-associated genes using a grey-blue scale. **D**, Representative immunofluorescence staining for Crystallin Alpha B (CRYAB, magenta) in cryo-section from a bulged region of a left carotid artery from a VSMC lineage-labeled Myh11-Confetti animal 5 days after ligation surgery (n=3 animals). Arrowheads point to CRYAB+Confetti+ cells and arrows indicate CRYAB-Confetti+ cells. Signals for CRYAB (magenta), Confetti proteins and DAPI (white) are shown as indicated. Green autofluorescence from the elastic laminae outlines the medial layer. Scalebar = 30 μm. **E**, UMAP of scRNA-seq dataset from VSMC-lineage-labeled cells from Myh11-Confetti-Apoe animals fed a high fat diet (Dobnikar et al., 2018). Left panel show summarized expression of genes with Path1-induced expression in the mouse D5 dataset (Path1 gene clades 1, 2, 5, 7, 8, 10) on a scale from blue (low) to red (high). Right panels show expression of VSMC-derived cell markers (*Myh11*: contractile, *Ly6a*: mesenchymal stem cell-like; *Chad*: osteochondrogenic; *Cd68*: macrophage) in grey-blue scales. **F**, Immunofluorescence staining for FBLN2 (magenta) in arterial cryo-sections from VSMC-lineage-labeled Myh11-EYFP-Apoe animals fed a high fat diet for 24 weeks. The left panel shows a merge with signals for the EYFP VSMC-lineage label (yellow). Green autofluorescence from the elastic laminae outlines the medial layer. Arrows point to EYFP+FBLN2+ cells. P: plaque, M: Medial layer, L: Lumen. Scalebar = 100 μm. **G**, Immunohistochemistry co-staining for FBLN2 (brown) and αSMA (blue) in FFPE sections from human carotid artery plaques. Representative of samples from 4 patients. Scalebar = 200 μm.

To evaluate the relevance of proliferation-associated, injury-induced genes in atherosclerosis, we assessed the expression of Path1-induced genes in VSMC-derived plaque cells^15^. This demonstrated anti-correlation with *Myh11* levels and overlap between the injury-response genes and markers of phenotypically modulated VSMCs in lesions (*Chad, Ly6a*, Figure 5E). Additionally, genes showing increased expression along the proliferation-associated Path1 included factors previously associated with vascular disease, such as *Lum* and *Tnfrsf11b*^15-17,32-34^. We also detected FBLN2-expression (a Path1-induced gene) in a subset of VSMC-derived plaques cells in Myh11-Confetti-Apoe animals, in particular in the lesion core (Figure 5F). Importantly, FBLN2 was detected in *α*SMA-stained cells in human carotid artery plaques (Figure 5G), indicating that this signature is also relevant for human disease.

Taken together, we identify two related but distinct injury-responses in VSMCs. Interestingly, Path2 is mainly characterized by enrichment for genes associated with protein folding, which has been linked to cholesterol responses^35^. In contrast, genes defining Path1, which represents a transition in cellular state associated with injury-induced VSMC proliferation, are also expressed in human atherosclerosis.

### SCA1 expression marks “first responder” VSMCs with increased proliferative capacity

We previously identified a small subset of SCA1-expressing VSMCs in healthy arteries, which express a “Response Signature” suggestive of cell activation^15^. The frequency of SCA1+ cells increase in vascular disease^15,16,36^ and the kinetics of SCA1 induction after injury correlates with emergence of KI67+ cells (Figure 6A, Figure 1E). Further indicating an association between SCA1 expression and VSMC proliferation, the *Ly6a*/SCA1 transcript was detected in Path1-associated cells in the D5 scRNA-seq dataset and *Ly6a*/SCA1+ cells were juxtaposed to *Mki67*+ cells, preceding those in pseudotime (Figure 6B). Interestingly, the “Response Signature” expressed by SCA1-positive VSMCs in healthy arteries^15^ was also induced in Path1-specific cells (Figure 6B), suggesting that SCA1 expression may mark VSMC that have undergone partial transition towards a proliferative state.

**Figure 6:**
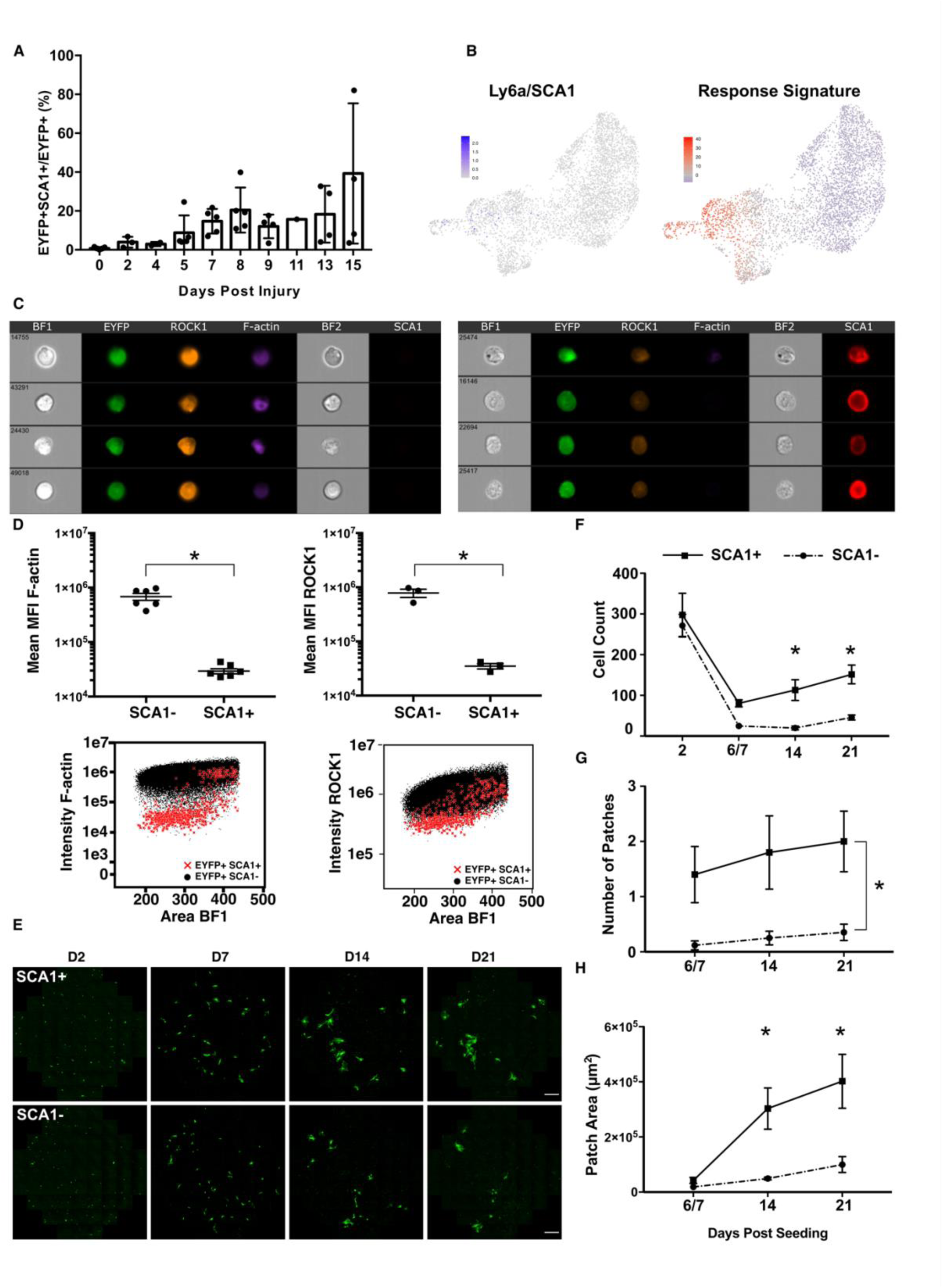
SCA1 marks VSMCs with increased proliferative capacity. **A**, Percentage of EYFP+ VSMCs expressing SCA1 in left carotid arteries of Myh11-EYFP-Ki67/RFP animals analyzed by FACS, showing mean ± standard deviation at indicated timepoints after ligation (total 32). **B**, UMAP of D5 scRNA-seq dataset showing expression of Ly6a/SCA1 using a scale from grey-blue (left) or summarized expression of the “Response Signature” identified in SCA1+ VSMCs from healthy arteries (Dobnikar et al., 2018) on a blue-red scale (right). **C**, ImageStream bright field (BF) and fluorescence images of medial cells from the aorta of VSMC lineage-labeled Myh11-EYFP animals stained for SCA1 (red), ROCK1 (orange) and F-actin (magenta, phalloidin) with VSMC-lineage label (EYFP) in green. **D**, Quantification of mean fluorescence intensity (MFI, top panels) and dot plots showing intensity (lower panels) for F-Actin (left) and ROCK1 (right) in EYFP+SCA1+ (red crosses) and EYFP+SCA1- (black circles) cells gated as shown in Figure VI of the Online-only Data Supplement. Cells from 3 animals were analyzed separately, indicated by dots in top panels, that also show mean and S.E.M. Asterisks indicate P<0.05, t-test. **E**, Live cell confocal images of 500 EYFP+SCA1+ (top) or EYFP+SCA1- (lower) cells from Myh11-EYFP animals mixed with 4500 medial cells from wild type animals in a 96-well plate. Cells were imaged at indicated timepoints using Opera Phenix system. EYFP signals are shown in green. Scalebar = 100 μm. **F-H**, Quantification of cell count per well (F), number of patches per well (G) and mean area of patches per well (H) for wells containing EYPF+SCA1+ (squares, solid lines) and EYFP+SCA1-cells (circles, dotted lines) at indicated timepoints post seeding. Plots show mean and S.E.M of cells from three animals analyzed separately. Statistical significance of cell count differences at individual days was tested by ANOVA (F); patch numbers were fitted to a generalized linear model (G); multiple linear regression was performed of log transformed data for patch area (H). Asterisks indicate P<0.05.

To test whether functional differences are associated with the SCA1-expression in healthy arteries, we analyzed FACS-isolated SCA1-positive and SCA1-negative lineage-labeled EYFP+ VSMCs from non-injured Myh11-EYFP animals. SCA1-positive VSMCs had remarkably reduced F-actin levels, demonstrated by significantly reduced phalloidin staining compared to the SCA1-negative counterparts (Figure 6C, D, Figure VI in the Online-only Data Supplement). Reduced F-actin in SCA1+ cells was accompanied by significantly lower levels of ROCK1, consistent with reduced *Rock1* transcript levels along Path1 during the injury response (Figure 5C). To compare cell proliferation, we adapted the *in vitro* clonal proliferation assay; mixing 500 SCA1+ or SCA1-EYFP+ cells from aortas of healthy, VSMC-lineage traced animals with medial cells from wild-type animals and periodic live imaging over 3 weeks of culture (Figure 6E). There was no difference in cell numbers 2 days after seeding, demonstrating equal survival. However after 1 week of culture, significantly more cells were detected in SCA1+ compared to SCA1-samples and this difference persisted throughout the experiment (Figure 6E, F). The increasing cell number resulted from emergence of coherent patches of EYFP+ cells (Figure 6E). In SCA1+ samples, 1-3 patches were observed per well 1 week after seeding (2.4 patches per well on average). Patch number remained approximately constant, whereas the size of individual patches increased over time (Figure 6G, H). In contrast, wells containing SCA1-EYFP+ cells did not contain patches after 1 week of culture (Figure 6G) and patches were observed at low frequency in SCA1-cultures at later timepoints (4/18 wells).

This analysis demonstrates that SCA1-expressing cells in healthy vessels are phenotypically and functionally distinct from the bulk of VSMCs. The 8-fold increased patch number and faster kinetics of clone formation for SCA1+ VSMCs suggests that these cells might act as “first responder” cells in healthy arteries exposed to disease-inducing stimuli.

### Evidence for priming of VSMCs in human arteries

SCA1 does not have an obvious human orthologue, preventing direct translation to human disease. Therefore, to assess whether healthy human arteries contain similarly primed cells, we performed scRNA-seq of cells from the medial layer of a histologically normal human aorta. Cells formed a single population that was split into 4 clusters without clearly defined borders (Figure 7A). Most cells expressed a contractile *MYH11*+ signature, consistent with the absence of signs of vascular disease, but reduced levels of contractile genes were observed in cell cluster 3 and a subset of cells in cluster 0. Anti-correlating with *MYH11*, gradually higher expression of *COL8A1, MGP* and other synthetic genes was detected through clusters 2, 0 and 3, suggesting that different extents of phenotypic switching exist in human vessels (Figure 7A). Transcripts for orthologues of injury-induced proliferation-associated genes, including *FBLN2* and *LUM*, were also detected in Cluster 0 and 3 (Figure 7A). To assess whether the healthy human aorta contains cells corresponding to those we identified in mouse vessels, we first tested whether genes associated with Path1 in the mouse D5 dataset show differential expression in cluster 0 versus cluster 1 (Figure 7B). Path1 downregulated genes generally showed lower expression in cluster 0 versus cluster 1, whereas most Path1-upregulated genes were detected at higher levels in Cluster 0. The “Response Signature” defining SCA1+ VSMCs in healthy mouse arteries^15^ also showed pronounced differences across the human VSMC dataset, with low levels in cluster 1 and high expression in some cells from cluster 0 and 3 (Figure 7C). This suggests heterogeneity of VSMC in the medial layer of human aorta, with phenotypic modulation similar to that found in SCA1+ mouse VSMCs in healthy arteries and after vascular injury. Interestingly, only few genes from injury Path1-associated gene clade 2, that included most cell-cycle genes, showed differential expression in the human dataset, consistent with the notion that VSMCs in healthy arteries are largely quiescent (Figure 7B, yellow dots).

**Figure 7:**
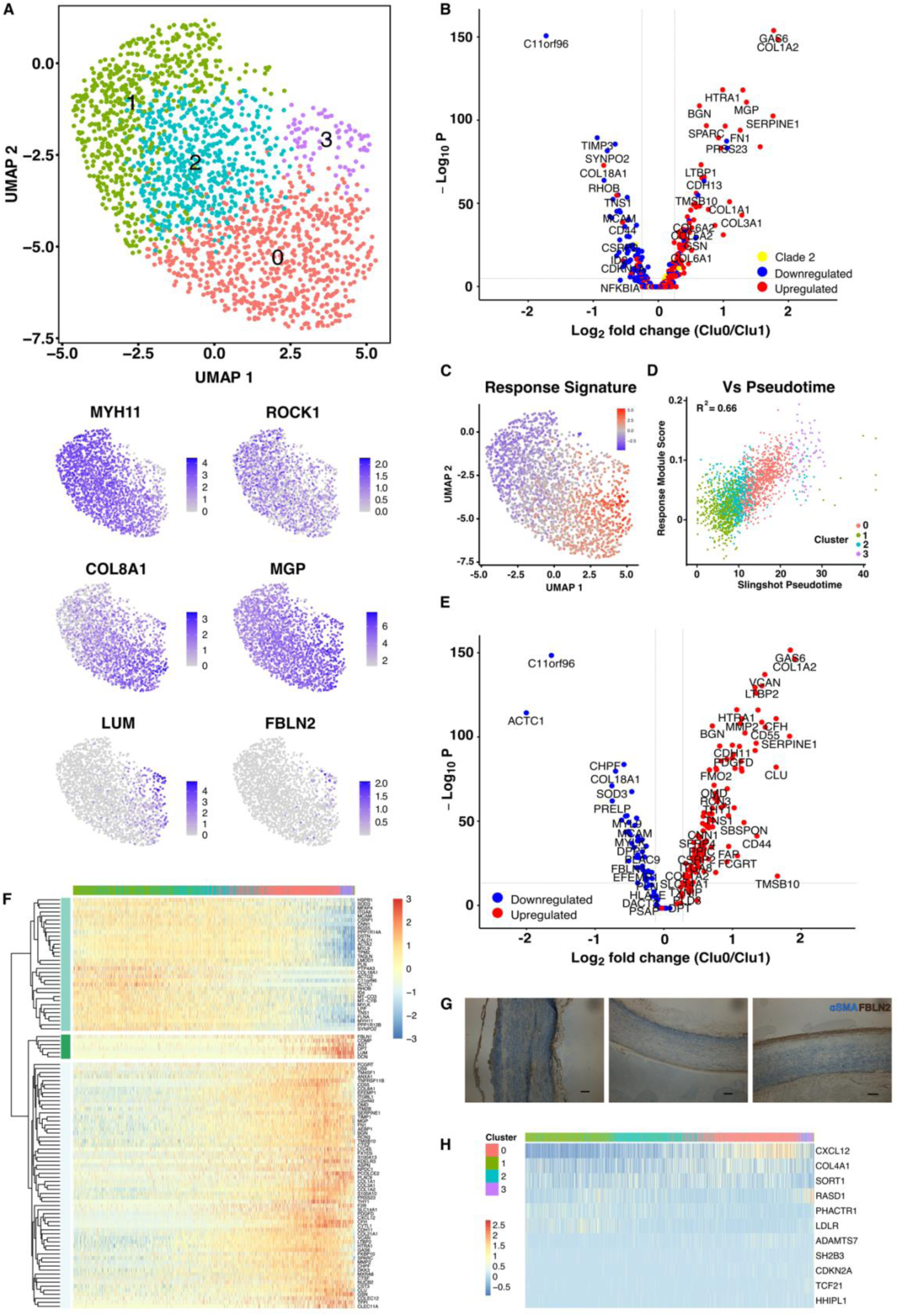
A subpopulation of human VSMCs in healthy arteries expressing a primed gene signature. **A**, Uniform Manifold Approximation and Projection (UMAP) of 10X chromium dataset of 1978 medial cells from a healthy human aorta showing cluster map (top) and expression level of marker genes using a grey-blue scale. **B**, Volcano plot showing differential expression between cluster 0 and cluster 1 cells. Only genes showing Path1-induced (red, Path1 gene clades 1, 5, 7, 8, 10), Path1-down-regulated (blue, Path1 gene clades 3, 4, 9) or cell-cycle-associated (yellow, Path1 gene clade 2) expression in the mouse D5 dataset are shown. **C**, UMAP of human dataset showing the summarized expression of the “Response Signature” identified in SCA1+ VSMCs from healthy arteries (Dobnikar et al., 2018), shown on human UMAP in a bluered scale. **D**, Scatter plot showing Module Score for the Response Signature versus the trajectory-inferred pseudotime for human scRNA-seq dataset, correlation coefficient 0.66. Cells are color-coded by cluster identity. **E**, Volcano plot showing differential expression between cluster 0 and cluster 1 cells. Only the top 200 genes with significant pseudotime-dependent expression that are expressed in both clusters (170 genes total, blue = pseudotime-downregulated genes; red = pseudotime-upregulated). **F**, Heatmap showing expression levels (on a scale from blue to red) of the top 100 genes with pseudotime-dependent gene expression (p-adj<0.005). Cells are ordered by pseudotime and top bar indicates cell cluster affiliation. Genes are clustered by Pearson correlation into 3 gene clades. **G**, Immunohistochemistry co-staining for FBLN2 (brown) and αSMA (blue) in FFPE section from healthy human aorta. Representative of six patients. Scalebar = 200 μm. **H**, Heatmap showing expression of the 11 genes included in the 27-SNP genetic risk score for coronary artery disease events (Mega et al., 2015), which show significant pseudotime-associated expression (p-adj<0.005). Cells are ordered by pseudotime and top bar indicates cell cluster affiliation.

The analysis above suggested that medial cells in human arteries are defined by a phenotypic spectrum, similar to what has been suggested^29^. To identify genes defining this spectrum in an unbiased manner, trajectory-inference was used to generate a pseudotime axis (Figure VII in the Online-only Data Supplement), which showed strong correlation with the Response Signature (R^2^=0.66, Figure 7D). As expected, pseudotime-dependent genes with reduced expression along the trajectory were detected at lower levels in cluster 0 compared to 1 (Figure 7E, blue dots), and were enriched for “muscle contraction” and “actin-binding” GO-terms (Table VII in the Online-only Data Supplement). Trajectory-induced genes showed increased expression in cluster 0 vs cluster 1 (Figure 7E, red dots) and were generally restricted to cells in cluster 3 and a subset of cluster 0 cells (Figure 7F). Enrichment for GO-terms related to extracellular matrix modification, cell adhesion, tissue development and response to TGF-beta (Figure VIII and Table VII in the Online-only Data Supplement) suggested that a VSMCs in healthy human vessels displaying evidence of phenotypic activation. In line with this idea, pseudotime-induced genes included growth factor binding proteins (*LTBP2, HTRA1*) and *CXCL12* that encodes stromal cell-derived factor 1 (SDF1) and is linked genetically to cardiovascular disease^37^. This analysis suggested that human arteries contain a subpopulation of VSMCs in a state corresponding to that defined by SCA1 expression in mouse. To verify this idea and add positional information for such primed VSMCs, we stained healthy human aorta sections for FBLN2, which is also pseudotime-induced in the human dataset (Figure 7F and Table VII in the Online-only Data Supplement). FBLN2+ cells were detected in the medial layer (Figure 7G) – in addition to adventital and endothelial staining – but at lower frequency compared to in lesions (Figure 5G). FBLN2+ medial cells did not cluster to specific regions that could represent pre-atheromas not detected in histological examination, instead, we find that FBLN2+ cells are dispersed in the medial layer of human arteries.

Interestingly, of the 23 genes associated with the SNP-27 coronary artery disease risk score panel^38^ that were included in our dataset, 11 showed differential expression along the trajectory; including *TCF21, CXCL12* and *ADAMTS7* (p<0.005; Figure 7H). This statistically significant enrichment strongly indicates that the changes along the pseudotime axis are relevant for human cardiovascular disease.

## DISCUSSION

Using an acute model of VSMC proliferation, we demonstrate that oligoclonal VSMC contribution to lesion formation results from selective activation of proliferation in very few pre-existing VSMCs and provide evidence that this mechanism is shared at early stages of atherosclerotic plaque formation. Immediately after injury, VSMCs form a continuous spectrum of phenotypes where a pseudotime axis connects quiescent cells with a contractile signature to proliferative cells. Cells along this trajectory include SCA1-expressing cells that share characteristics of the atypical VSMC we previously identified in healthy arteries^15^. The increased proliferative capacity observed for SCA1+ VSMCs further supports the notion that these cells are predisposed to react to activating signals. The demonstration that healthy human arteries also display transcriptional heterogeneity for genes linked genetically to cardiovascular disease, and contain cells displaying significant similarities to the SCA1+ cells in mouse arteries, suggests that VSMC priming could also underlie vascular pathologies in humans.

In addition to the pseudotime axis resulting in cell proliferation (Path1), we find evidence for another VSMC response at early timepoints after injury (Path2). Both response trajectories show increased expression of ECM-related factors and reduced contractile gene expression suggesting that both represent induction of a “synthetic state”. Increased expression of structural components and regulators of the ECM is consistent with the observation that VSMC proliferation is observed in arterial segments showing pronounced remodeling of the vessel wall (Figure 2). Yet, only a fraction of VSMCs within the arterial “bulges” exit quiescence to initiate proliferation and, while substantial VSMC loss is observed, most cells within these remodeled arterial segments remain as singlets. This selective VSMC activation and oligoclonality of lesional VSMCs appears at odds with the gradually changing cells states observed in scRNA-seq where SCA1 expressing cells partially overlap proliferating cells. We suggest that, rather than being a dedicated progenitor population, some cells are predisposed, or primed, for proliferation. In accordance with this idea, SCA1+ VSMCs show an 8-fold increased proliferation frequency *in vitro*, although proliferation was also observed in SCA1-cells, albeit with slower kinetics. We speculate that formation of patches in SCA1-sorted samples result from the induction of a Sca1 signature previously seen in cultured VSMCs^15^. An alternative idea is that activation of VSMC proliferation induce negative feedback mechanisms to prevent neighboring cells from exiting quiescence akin to lateral inhibition. Experimental testing of these ideas is not trivial. Firstly, SCA1 is expressed by other cell types in the vasculature, necessitating a dual lineage labeling approach^16,39^. Secondly, current SCA1-Cre drivers are not sufficiently highly expressed in medial cells to yield recombination-induced cell labeling^40^, probably due to the relatively lower expression level of *Ly6a* transcripts in VSMCs compared to, for example, adventitial and endothelial cells^15^.

The transcriptional signature defined by the contractile-to-proliferative axis in post-injury VSMCs shares considerable overlap with transcriptional states of VSMC-derived cells in other vascular disease models, including atherosclerosis and aneurysm^15-17,32-34^. We therefore propose that the mechanisms acting at the onset of VSMC proliferation after injury also regulate early steps of atherosclerotic plaque development. Consistently, we observe clonal VSMC contribution at early stages of plaque development, even before formation of the fibrous cap, in contrast to a study suggesting that VSMC investment results from migration along the fibrous cap and that VSMC-derived cells in the plaque core are derived from expanding aSMA+ cells^10^. Despite these apparent discrepancies, our findings are in accordance with the idea of a phenotypically modulated, plastic cell state that underpins atherosclerotic lesion VSMC infiltration^15^. Such a state has been defined by expression of *Lgals3*, which is present prior to cap formation^32^ and also SCA1^16,17,33^, consistent with the observation of proliferation-associated SCA1+ cells in our dataset. We did not observe VSMC-derived cells expressing an osteochondrocytic signature present in atherosclerotic lesions^15,33^. Whether this is due to model-specific differences or the time point of analysis remains to be determined. However, we note that the osteochondrocytic phenotype is more pronounced at late time points and was not observed in studies limited to early-mid stage disease^32,33^.

Our study identifies additional phases of activation where VSMCs that could be subject to regulation, including VSMC priming, cell cycle activation, VSMC loss and migration across the intimal layer. Understanding how documented regulators of VSMC function in disease^1,36,41^ and novel pathways - such as retinoic acid signaling and efferocytosis identified by scRNA-seq analysis of atherosclerotic plaque cells^16,33^ -impact on these mechanisms will provide important insight into how targeting of vessel wall cells could be achieved to limit cell accumulation and disease severity. The existence of cells in human arteries that correspond to the murine SCA1+ VSMCs and the genetic evidence linking variable expression to cardiovascular disease highlight this cell population a promising starting point.

## Supporting information

Online-only Data Supplement

## Acknowledgments

The authors would like to thank the Wellcome Trust-Medical Research Council, Institute of Metabolic Science, Metabolic Research Laboratories, Imaging core, Wellcome Trust Major Award [208363/Z/17/Z] for assistance with confocal imaging, the National Institute for Health Research Cambridge Biomedical Research Centre Cell Phenotyping Hub for cell sorting, Katarzyna Kania and colleagues (Genomics Core Facility, Cancer Research UK Cambridge Research Institute) and Kristina Tabbada and colleagues (Babraham Institute Sequencing Facility, funded by the UKRI-BBSRC Core Capability Grant) for 10x chromium single cell library preparation and Illumina sequencing, Simon Andrews and Felix Krueger at the Babraham Institute’s Bioinformatics facility for raw data processing and read alignment and all members of the Jørgensen team for helpful discussions.

## Disclosures

None

**Online-only Data Supplement includes Full Methods, Figures I-VIII, Tables I-VII**

