## Supplementary material for "Primed smooth muscle cells acting as first responder cells in disease": Online-only Data Supplement

Worssam/Lambert/Oc *et al.*

#### Content of this document

Extended Methods

Supplemental figures

- Figure I: Examples and quantification of whole-mounted left carotid arteries of VSMC-lineage labeled Myh11-Confetti animals
- Figure II: Tissue explant analysis
- Figure III: Feature plots of mouse D7 scRNA-seq dataset including cell cluster 11
- Figure IV: Trajectory inference for mouse D5 scRNA-seq data
- Figure V: Gene ontology (GO) analysis of genes showing trajectory-induced expression in mouse D5 scRNA-seq dataset
- Figure VI: ImageStream analysis
- Figure VII: Trajectory inference for human VSMC scRNA-seq dataset
- Figure VIII: Enriched gene ontology terms for human pseudotime-associated genes

References

#### Separate files

Supplemental tables

- Table I: Antibodies used in this study
- Table II: Carotid arteries analyzed by whole mount imaging
- Table III: Top marker genes for cell clusters in mouse D7 scRNA-seq data
- Table IV: Genes showing significant association with Path1 and Path2 in mouse D5 scRNA-seq data
- Table V: Gene ontology analysis of genes showing induced expression along Path1 and Path2 in mouse D5 scRNA-seq dataset
- Table VI: Genes showing differential expression in human scRNA-seq data
- Table VII: Gene ontology analysis of genes of genes showing pseudotime-associated gene expression in human scRNA-seq data

### EXTENDED METHODS

##### *Human tissue*

Anonymized human arteries (plaques and normal aortas) were obtained from patients undergoing carotid endarterectomy or coronary artery bypass/valve replacement, respectively, under informed consent using protocols approved by the Cambridge or Huntingdon Research Ethical Committee.

##### *Animals and procedures*

Animal experiments were approved by the local ethics committee and were performed according to UK Home Office regulation under project license P452C4595. All alleles have been described previously; Myh11-CreERt2 is a Y-linked transgene that confers expression of a tamoxifen-inducible Cre recombinase in smooth muscle cells (Chakraborty et al., 2019; Wirth et al., 2008), Rosa26-Confetti (Snippert et al., 2010) and Rosa26-EYFP (Srinivas et al., 2001) are Cre-recombination reporter alleles, Ki67-RFP is an insertion in the Mki67 locus resulting in expression of a KI67-RFP fusion protein (Basak et al., 2014) and the mutant Apoe allele sensitizes mice to high fat diet (HFD)-induced atherosclerosis development (Piedrahita et al., 1992). Unless indicated, VSMC lineage labeling was done by 10 intraperitoneal injection of 0.1 mg tamoxifen (1 mg/mL in corn oil) over 2 weeks into Myh11-CreERt2, Rosa26-Confetti (Myh11-Confetti), Myh11-CreERt2, Rosa26-Confetti, $Apoe^{-/-}$ (Myh11-Confetti-Apoe), Myh11-CreERt2, Rosa26-EYFP (Myh11-EYFP), Myh11-CreERt2, Rosa26-EYFP, $Apoe^{-/-}$ (Myh11-EYFP-Apoe) and Myh11-CreERt2, Rosa26-EYFP, Ki67-RFP (Myh11-EYFP-Ki67/RFP) animals. All animals were rested for at least 1 week to allow tamoxifen washout. The Myh11-CreERt2 is Y-linked, so all VSMC lineage-tracing experiments were performed using males. Carotid ligation was performed as previously described (Chappell et al., 2016). Briefly, animals were given pre-operative analgesic (~0.1 mg/kg body weight, Buprenorphine) subcutaneously, anaesthetized with 2.5-3% isoflurane by inhalation (1.5 L/min) and the left carotid artery was tied off with a silk suture. Animals were culled 2-28 days after surgery by CO2 asphyxiation and perfused with cold phosphate buffered saline (PBS) before tissue removal. Lineage-labeled Myh11-Confetti-Apoe animals were fed a HFD (Special Diets Services, containing 21% fat and 0.2% cholesterol) for 9-24 weeks as indicated, starting 1 week after the last tamoxifen injection.

##### *Tissue processing for imaging*

Mouse arteries were dissected free of any adipose tissue and fixed in fresh 4% formaldehyde (Sigma) for 20 minutes at room temperature. Carotid arteries were stained with 4', 6-diamidino-2-phenylindole (DAPI, 1 µg/mL in PBS) overnight at 4 °C, cleared using RapiClear 1.52 (Sunjin Lab) for at least 5 hours and mounted in RapiClear 1.52 using iSpacers (Sunjin Lab) for imaging. Fixed arteries from Myh11-Confetti-Apoe, Myh11-EYFP-Apoe animals, and post-imaging carotid arteries, were frozen in optimal cutting temperature (OCT, Tissue-Tek) compound after cryopreservation in 30% (w/v) sucrose in PBS an equilibration in sucrose:OCT (1:1). Sections were cut on a cryostat microtome onto Superfrost™ Ultra (thin, 14 µm) or Superfrost™ Ultra Plus Slides (thick, 100 µm)(Thermo scientific) and stained with DAPI (1 µg/mL in PBS) for at room temperature before mounting in RapiClear 1.52.

Human arteries were formaldehyde fixed and paraffin embedded (FFPE) and 4 µm sections cut onto Superfrost™ Ultra Plus slides.

##### *Tissue explants*

Aortas were isolated from VSMC-lineage labeled animals (Myh11-Confetti or Myh11-EYFP), dissected free of perivascular adipose tissue and pre-digested in Opti-MEM (Gibco) supplemented with 1 mg/mL Collagenase IV (Invitrogen) and 1 U/mL Elastase (Worthington) for 10 minutes at 37˚C to allow removal of the adventitial layer. Arteries were kept in Opti-

MEM overnight and 1 $mm^2$ tissue sections cut with minimal mechanical manipulation. Explants were pinched with sharp forceps to create an internal injury site and embedded in Matrigel in an 8-well chamber slide (Ibidi). Growth medium (Opti-MEM supplemented with 10% (v/v) fetal bovine serum (FBS), 100 U/mL penicillin, 100 mg/mL streptomycin and 20 ng/mL PDGF was added and changed every 2-3 days. Explants were fixed directly (day 0) or after 8 days of culture in 2% (w/v) methanol-free paraformaldehyde (Thermo Fisher) and 0.5% (w/v) glutaraldehyde (Sigma Aldrich) in PBS for 30 minutes at 4 °C. After washing in PBS, explants were mounted in RapiClear 1.47 (Sunjin Lab) using iSpacers (Sunjin Lab).

***Immunostaining of tissue sections***
Cryosections were thawed, rinsed in PBS and permeabilized for 20 minutes in 0.5% (v/v) Triton X-100 (Sigma Aldrich) in PBS at room temperature. Sections were blocked for 1 hour at room temperature in 1% (w/v) bovine serum albumin and 10% (v/v) of either normal goat serum (Dako), or normal donkey serum (Abcam) in PBS and incubated with 2.5 µg/mL anti-FBLN2 (Abcam, ab251662), 1 µg/mL anti-CRYAB (Abcam, ab13497) or isotype controls (rabbit IgG Abcam, ab37415), diluted in blocking buffer overnight at 4°C or 30 minutes at 37˚C. Primary antibodies were removed by washing 3x 5 minutes in PBS before incubation with Alexa-647 conjugated secondary antibodies, diluted in blocking buffer, for 1 hour at room temperature. Sections were washed 2x 5 minutes in PBS and nuclei stained with DAPI (1 µg/mL) for 10 minutes at room temperature, before rinsing in PBS and mounting in RapiClear 1.52 (Sunjin Lab).

Formaldehyde fixed paraffin embedded (FFPE) sections were dewaxed in Xylene, rinsed in water and antigen retrieval performed in Citrate-based Antigen Retrieval Solution (pH6, Vector Labs) for 30 minutes in the Aptum 2100 Retriever Pressure Cooker. Blocking was done by first incubating in Peroxidase Blocking Solution (Abcam, 10 minutes), washing 3x 5 minutes in PBS and then incubating with blocking in antibody diluent (SignalStain, 10 minutes). Blocked sections were incubated with anti-FBLN2 (Abcam, ab251662) or isotype control (Abcam, ab37415) in antibody diluent overnight at 4°C, washed 3x in PBS, stained with HRP-conjugated anti-Rabbit (Cell Signaling Technology, 8114) in antibody diluent for 30 minutes at room temperature, washed 3x in PBS and HRP visualized using DAB peroxidase substrate (SignalStain). After rinsing in water, sections were stained with anti-αSMA (DAKO, M0851) in Superblock diluent (Thermo Fisher) for 1 hour at room temperature, washed 3x in PBS and incubated with biotin-coupled anti-Mouse (DAKO, E0433) in Superblock diluent for 30 minutes at room temperature, washed 3x in PBS and incubated in Vectastain avidin-coupled alkaline phosphatase (AP) reagent (Vector Labs) for 30 minutes, before rinsing in PBS and incubating with Blue AP substrate solution (Vector Labs) for 20 minutes, all at room temperature. Finally, sections were washed in PBS and mounted in VectaMount mounting media (Vector Labs).

***Confocal microscopy and image analysis***
Confocal imaging was done with an SP8 laser scanning microscope (Leica). Sequential, resonant scan mode with laser lines and detectors set for maximal sensitivity without spectral overlap for DAPI (405 laser, 417-508 nm), CFP (458 laser, 454-502 nm), GFP (488 laser, 498-506 nm), YFP (514 laser, 525-560 nm), RFP (561 laser, 565-650 nm) and Alexa Fluor 647 (633 laser, 650-700 nm) and oil immersion objectives (20x, 40x) were used. Imaging was done in tile scan mode and the distance between sections in Z-stacks was 5-6 µm for whole mounted carotid arteries, 5 µm for tissue explants, 3-4 µm for thick cryosections and 3 µm for cryosections. Z-compensation (laser power) was used to normalize fluorescent protein intensity throughout the tissue in thick specimens. Data was acquired at an optical section resolution of 1024 x 1024 and tiles were stitched using the mosaic merge function in LASX software (Leica). Image analysis was done using Imaris software (9.0.2) to adjust brightness and contrast, generate maximal projections, for virtual cross sectioning, surface rendering and measurements.

In ligated carotid arteries, regions containing ≥3 adjacent cells of the same color were scored as a medial patch if the distance between the edges of neighboring cells was less than 0.5 µm (CFP), 1.0 µm (RFP, YFP) or 15 µm (GFP) and did not cross an elastic laminal layer. Patch size was scored as small (3-10 cells), medium (11-49 cells) or large (>50 cells) manually. Bulged regions were scored as continuous stretches where the outer elastic lamina diameter ≥450 µm. Intimal patches were scored as contiguous regions with ≥5 cells of the same color inside of the inner elastic lamina. The lengths of the bulged regions were measured as the length of the outer elastic lamina. VSMC death was measured by outlining acellular (DAPI-Confetti-) medial areas in virtual cross sections using the Measurement Points tool in Imaris. For each artery, the percentage acellularity relative to total medial area was calculated for 5 sections in a region with normal diameter and 5 sections in a "bulged" region (if present). Arteries where image quality was affected by tissue damage or autofluorescence (8 arteries) or arteries analyzed more than 10 days after injury but did not show any sign of reaction to surgery (e.g. adventitial expansion, 3 arteries) were excluded from analysis.
To analyze initial VSMC contribution in early stage lesions, plaques containing <50 Confetti+ cells per section were included. For each lesion, the number of fluorescent proteins was determined and the position of Confetti+ cells within lesions scored.
For tissue explants, the surface of individual patches was rendered using default settings for surface area detail level and enabling background subtraction (a comparison of rendered and original explant images is shown in Figure II in the online data supplement. Rendered images were used to calculate the number and volume of patches that exceed 30,000 $\mu m^3$ for CFP/RFP/YFP or 5,000 $\mu m^3$ for GFP.

##### *Isolation of single cell VSMC suspensions*
Aortas or carotid arteries from wild type or VSMC lineage-labeled animals (Myh11-Confetti, Myh11-EYFP or Myh11-EYFP-Ki67) were cut open longitudinally and the endothelium removed by gentle scraping with a cotton bud before removal of the adventitia as described above. The medial layer was digested to a single cell suspension in DMEM supplemented with 2.5 mg/mL Collagenase IV (Invitrogen) and 2.5 U/mL Elastase (Worthington) at 37 ˚C.

##### *Flow cytometry assisted cell sorting (FACS), Flow cytometry, Image stream*
Single cell suspensions incubated with 5 µg/mL TruStain FcX anti-mouse CD16/32 antibody (Biolegend) in FACS buffer (0.5% (w/v) BSA in PBS) for 15 minutes on ice to block Fc-receptors, incubated with primary or isotype control antibody for 15 minutes at room temperature and washed twice in FACS buffer. Where needed, cells were incubated with secondary antibody in FACS buffer for 15 minutes at room temperature and washed twice in FACS buffer. Intracellular targets (ROCK1, Proteintech, 21850-1-AP) were stained using the Foxp3 staining buffer set (eBioscience) and phalloidin-iFluor™ 350 was included with secondary antibodies.
Stained cells were filtered (40 µm) and either sorted (BD FACSAria™ III, BD Bioscience) or subjected to either flow cytometry (Accuri C6 or BD Fortessa, BD Bioscience) or Imagestream (Amnis® ImageStream®XMk II, Luminex) analysis. Flow cytometry compensation was done using single stained samples and gates were defined based on samples stained with control antiserum and wild type cells.

##### *Clonal proliferation assay*
Medial cells from VSMC-lineage labeled Myh11-Confetti animals were mixed with medial cells from wild type animals in a 1:3 ratio and a total of 5,000 cells seeded per well of a 96-well imaging plate (CellCarrier-96 Ultra, Perkin Elmer) in DMEM supplemented with 10% (v/v) FBS, 100 U/mL penicillin, 100 mg/mL streptomycin and 2 mM Glutamine. Medial cells from lineage-labeled Myh11-EYFP animals stained for Sca1 and 500 Sca1+EYFP+ or Sca1-EYFP+ VSMCs were isolated by FACS and mixed with 4500 wild type medium cells before plating. Medium was changed every 2-3 days and the cells were imaged 4 days after plating and after 1, 2, 3 weeks of culture using an Opera Phenix high content screening system

(Perkin Elmer). Image analysis was done using Harmony software (Perkin Elmer) and quantification was performed in Fiji (Schindelin et al., 2012). Patches were defined as an area with three or more contiguous EYFP+ cells. To calculate the area of patches images were thresholded after enhancing local contrast (CLAHE) (ZUIDERVELD, 1994). The "analyze particles" function was then used to generate a mask of the image, and areas of pre-identified patches were extracted.

***scRNA-seq data generation***
Murine cells: Single cell suspensions of medial cells were generated from VSMC lineage-labeled Myh11-EYFP-Ki67/RFP animals as described above. For the D5 dataset, ligated carotid arteries from six animals were isolated 5 days after surgery and cells pooled prior to FACS isolation of EYFP+ cells, 20,000 EYFP+ cells were loaded onto the Chromium system (10x Genomics) and amplified cDNA libraries generated using the 3' Gene Expression v3.0 kit (10x Genomics) were sequenced using 150 cycle protocol (NovaSeq, Illumina). To generate the 10X D7 dataset, cells from the ligated left carotid arteries of five animals dissected 7 days after surgery were pooled and enriched for proliferating RFP+EYFP+ cells during FACS. A total of 100 EYFP+RFP+ cells were supplemented with 20,000 unselected EYFP+ cells before loading onto a 10X Chromium system. Amplified cDNA was generated using the 3' Gene Expression v2 kit and sequenced using a paired end protocol (HiSeq 4000, Illumina). The Smart-seq2 protocol (Picelli et al., 2014) was used to generate amplified cDNA libraries from EYFP+RFP+ and EYFP+RFP- cells isolated from the ligated arteries of animals 7 days after surgery and EYFP+ cells from unligated litter mate control animals in 2 separate experiments (4 or 5 ligated arteries and 1or 2 control arteries were pooled respectively in the two experiments). Cells were processed as described (Dobnikar et al., 2018) with addition of ERCC spike controls (diluted 1:80,000,000, Invitrogen). Pooled cDNA libraries were sequenced (HiSeq 2500, Illumina).

Human cells: Healthy human aorta from a 65-year-old male was dissected to remove the adventitial and intimal layers. The medial layer was cut into 1 $mm^2$ pieces, a single cell suspension was obtained by digestion in 3 mg/mL collagenase IV and 1 U/ml elastase. Following staining with 7-Aminoactinomycin D (7-AAD), cells were fixed in Lomant's Reagent (dithio-bis(succinimidyl propionate, DSP) at 4 °C overnight. Viable cells were isolated by FACS based on 7-AAD exclusion. Cells were de-crosslinked in dithiothreitol (DTT) incubation at 37°C for 30 minutes. A total of 20,000 cells were loaded onto a 10X Chromium system and amplified cDNA libraries generated using the 3' Gene Expression v3.0 kit (10x Genomics) were sequenced using 150 cycle paired end protocol (NovaSeq, Illumina).

***scRNA-seq data analysis***
For the mouse D7 Smart-seq2 data, reads were aligned to the GRCm38 mouse genome using TopHat aligner v2.1 (Trapnell et al. 2009) and Htseq-count v0.8 (Anders et al. 2015) was used to count the number of read alignments per gene in the resulting bam files (idattr = "gene_id" and stranded = "no"). Cells with <200,000 total reads, <1,000 genes detected and >30% of ERCC controls were excluded from analysis and genes with mean expression <1 were removed. The function computeSumFactors from the Bioconductor R package scran (Lun et al., 2016b) was used to compute the normalization factors for individual cells.

The 10X Genomics Cell Ranger pipeline v.2.1.1 was used to align raw sequencing reads to GRCm38 mouse genome and count the number of reads aligning to each gene for D7 dataset. Cell Ranger pipeline v.3.1.0 was used to align raw sequencing reads to a custom genome (based on the GRCm38 mouse genome, including the open reading frame encoding EYPF (Srinivas et al., 2001) for D5 dataset. For the human data set, raw sequences were aligned to GRCh38 using the Cell Ranger pipeline v.3.1.0. A previously published 10X scRNA-seq dataset of VSMC-lineage labeled plaque cells from Myh11-Confetti-Apoe animals fed a high fat diet for 14 or 18 weeks were obtained from the Gene

Expression Omnibus (GEO) repository (accession number: GSE117963) (Dobnikar et al., 2018).

Quality control (QC), normalization and gene expression analysis was performed using the CRAN R package Seurat v.3.1.2 (Butler et al., 2018; Stuart et al., 2019) in R v.3.6.2. Cells with <5,000 UMI counts, <2,000 genes detected and >5% mitochondrial reads were excluded for the D5 dataset and for the D7 dataset cells with <5,000 UMI counts, <2,000 genes detected and >6% mitochondrial reads were excluded. For the plaque dataset, cells with <5,000 or >20,000 UMI counts, <1,000 or >5,000 genes detected and >9% mitochondrial reads were excluded from the analysis. For the human dataset, cells with <35,000 UMI counts, <1,000 genes detected and >10% mitochondrial reads were excluded.

Normalization was performed using SCTransform (Hafemeister and Satija, 2019) v.0.2.1 implemented in Seurat (mouse datasets) or scran (Lun et al., 2016a)(human dataset). Ribosomal gene expression was regressed out in the human dataset before dimensionality reduction. Principal component analysis (PCA) was performed with 3,000 highly-variable genes (HVG). The first 27 (mouse D5), 25 (mouse D7), 30 (mouse plaque) or 15 (human) principal components (PCs) were used, and clustering was done at resolution 0.8 (mouse D5), 1.6 (mouse D7), 1.1 (mouse plaque) or 0.6 (human).

Marker genes in the cell clusters in the mouse D7 dataset were identified through differential expression testing which was performed via Wilcoxon rank sum test using FindAllMarkers function with the log fold-change threshold of 0.5 and adjusted p-value threshold of 0.05 and visualized using the DoHeatmap function in Seurat. Mouse D7 Smart-seq2 and D7 10x datasets were integrated using Seurat v.3.2.1 (Stuart et al., 2019). Prior to integration, genes expressed in less than 3 cells and cells with less than 200 detected genes and more than 15% mitochondrial reads were removed from the D7 Smart-seq2 dataset. Each dataset was log normalized, and 2,000 HVGs were identified in each prior to integration. PCA was performed following the integration of the datasets, and the first 30 PCs were used for the UMAP visualization of the integrated dataset.

Differential expression analysis for cluster 0 versus cluster 1 in the human scRNA-seq dataset was done using DEseq2 (Love et al., 2014).

Trajectory inference was performed using ScanPy partition-based graph abstraction (PAGA) in ScanPy (Wolf et al., 2019) or the Bioconductor R package slingshot v.1.4.0 (Street et al., 2018). For the mouse D5 dataset cluster 1 was used as the starting cluster. No starting cluster was chosen for analysis of the human dataset, but identical results were obtained when selecting cluster 1 as starting cluster. Significance of differential expression along pseudotime was tested through CRAN R package gam v.1.16.1 with an FDR-adjusted p-value threshold of 0.05. For the mouse D5 dataset, a magnitude threshold of 0.5 (log scale) was also applied. Genes showing differential expression along pseudotime were hierarchically clustered using CRAN R package pheatmap v.1.0.12 with the Pearson correlation distance and complete linkage method. The optimal number of gene clades was determined through the elbow method which was performed via CRAN R package factoextra v.1.0.7.

Gene ontology (GO) term analysis of mouse Path1 gene clusters was performed with the enrichGO and compareCluster functions implemented in Bioconductor R package clusterProfiler v.3.14.3 (Yu et al., 2012) by using genome-wide annotation for mouse with Bioconductor R package org.Mm.eg.db v.3.10.0. Benjamini-Hochberg method was chosen as the multiple testing correction method, and p-value and q-value thresholds were set as 0.05 in GO term analysis.

To identify enriched GO terms showing pseudotime-associated expression in the human scRNA-seq dataset, genes were ranked by significance (FDR-corrected p-value) and the 200 top genes grouped using Pearson correlation into gene clades as described above. Gene enrichment analysis was done using g:Profiler (Raudvere et al., 2019). Multiple testing correction was done using the tailor-made algorithm g:SCS with p-value threshold 0.05.

The summarized expression level of gene subsets was calculated through their first principal components across the cells in the respective dataset and the Module Score calculated using the AddModuleScore function in Seurat. The gene list of the VSMC response signature was obtained from Dobnikar et al. (Dobnikar et al., 2018). Path1-induced genes included gene clades 1, 2, 5, 7, 8, 10.

***Statistical analysis***
Statistical analysis was performed in R or GraphPad Prism v7. The Shapiro–Wilk test was used to ascertain normal distribution, and based on the normality of the data, equal variance assessed using Bartlett or Levine tests. Tests used to assess statistical significance were selected based on whether data met assumptions for normality and equal variances, and are indicated in figure legends. Local regression analysis was used to fit a LOESS curve of patch number. To assess statistical significance of SCA1 expression status in the clonal proliferation assay, a generalized linear model was fitted for patch number whereas multiple linear regression used for patch area, as the data showed equal variance and linearity and the residuals were approximately normally distributed.

***Data availability***
The scRNA-seq datasets generated in this study (mouse D5 10x, mouse D7 10x, mouse D7 Smart-seq2, human 10x) have been deposited to the Gene Expression Omnibus (GEO) repository, accession number: will be made available prior to publication. The scRNA-seq dataset from VSMC-lineage-labeled plaque cells from high fat diet fed Myh11-Confetti-Apoe animals is available from GEO, accession number: GSE117963. Scripts used for data analysis are available upon request.

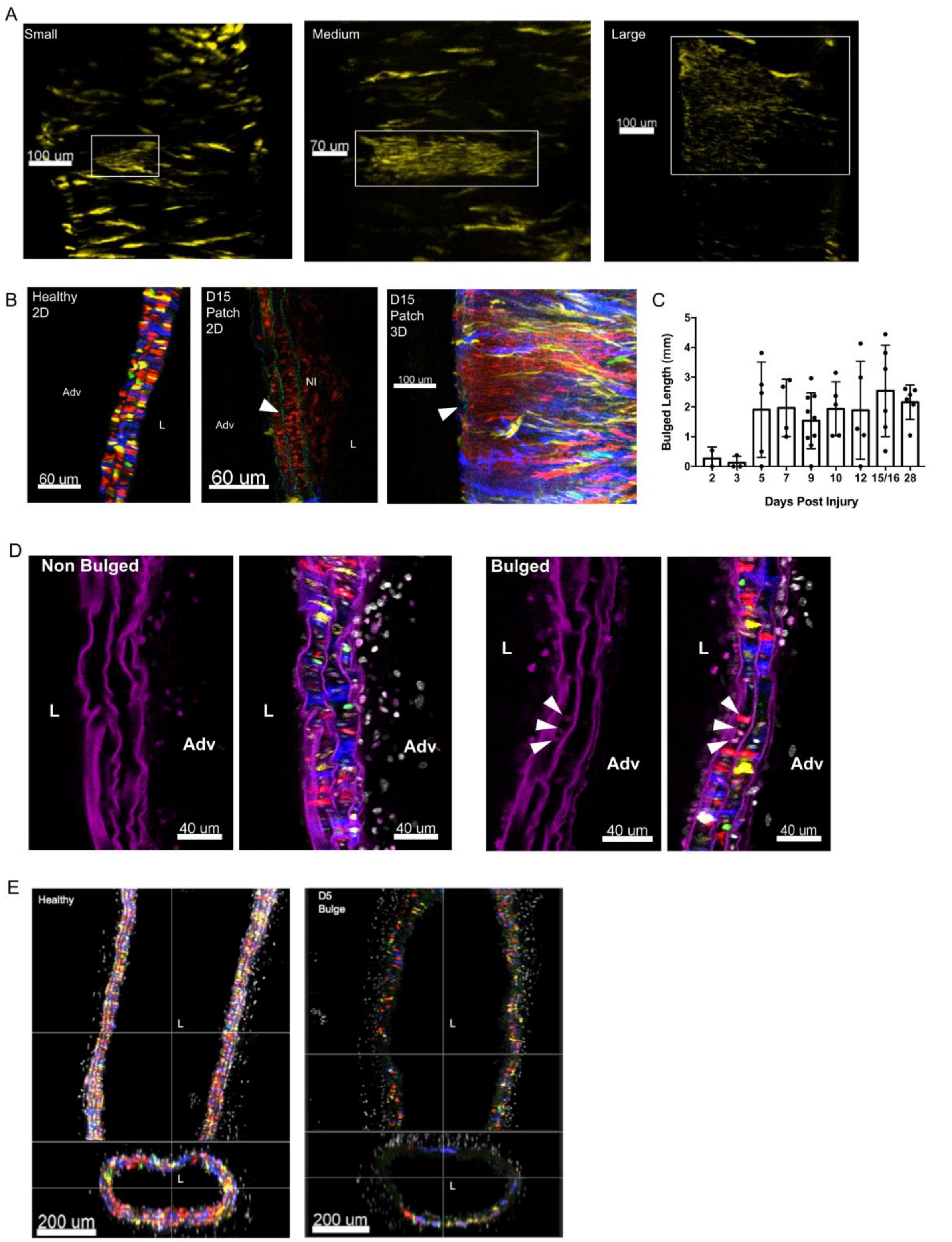


**Figure I: Examples and quantification of whole-mounted left carotid arteries of VSMC-lineage labeled Myh11-Confetti animals. A,** Confocal images (max projections) of ligated carotid arteries showing examples of small (left), medium (middle) and large VSMC patches (right). Only signals from the yellow fluorescent protein (YFP) of the Confetti reporter are shown. Scalebars = 100 μm (left, right), 70 μm (middle). **B,** Single Z-scans of carotid artery from unligated control animal (left) and ligated artery 15 days (D15) after surgery (middle) is shown to illustrate cellular disarray in regions with medial VSMC patches (RFP+, arrowhead). A neointimal (NI) RFP+ patch is also present. The adventitial (Adv) and luminal (L) sides are indicated. Scalebars = 60 μm. Right panel shows max projection of the D15 artery for comparison, with arrow head pointing to the RFP+ medial VSMC patch shown in the middle panel. Scalebar = 100 μm. **C,** Bar chart showing the number of neointimal patches as a fraction of medial patches for each artery at indicated time after surgery. Values for individual arteries, mean and S.E.M are shown. **D,** Representative confocal image (single Z-scan) showing EdU incorporation in a whole mounted carotid artery 5 days (5) after ligation (n=4 animals). The animal was injected with EdU intraperitoneally 16h prior to sacrifice. The two left panels show a non-bulged region and the two right panels show a "bulged" region from the same artery. EdU signal (magenta) is shown alone or merged with Confetti reporter and DAPI signals. The adventitial (Adv) and luminal (L) sides are indicated. Arrowheads point to three adjacent EdU+RFP+ cells in a medial patch. Scalebar = 40 μm. Note that substantial autofluorescence from the elastic lamellae is present in the magenta channel. **E,** Virtual horizontal (top) and transverse (lower) cross-sections generated in Imaris from whole mount images of carotid arteries from an unligated control animal (left) and a bulged region from a ligated artery 5 days (D5) after surgery (right). Signals for DAPI and the Confetti reporter are shown. Scalebars = 200 μm.

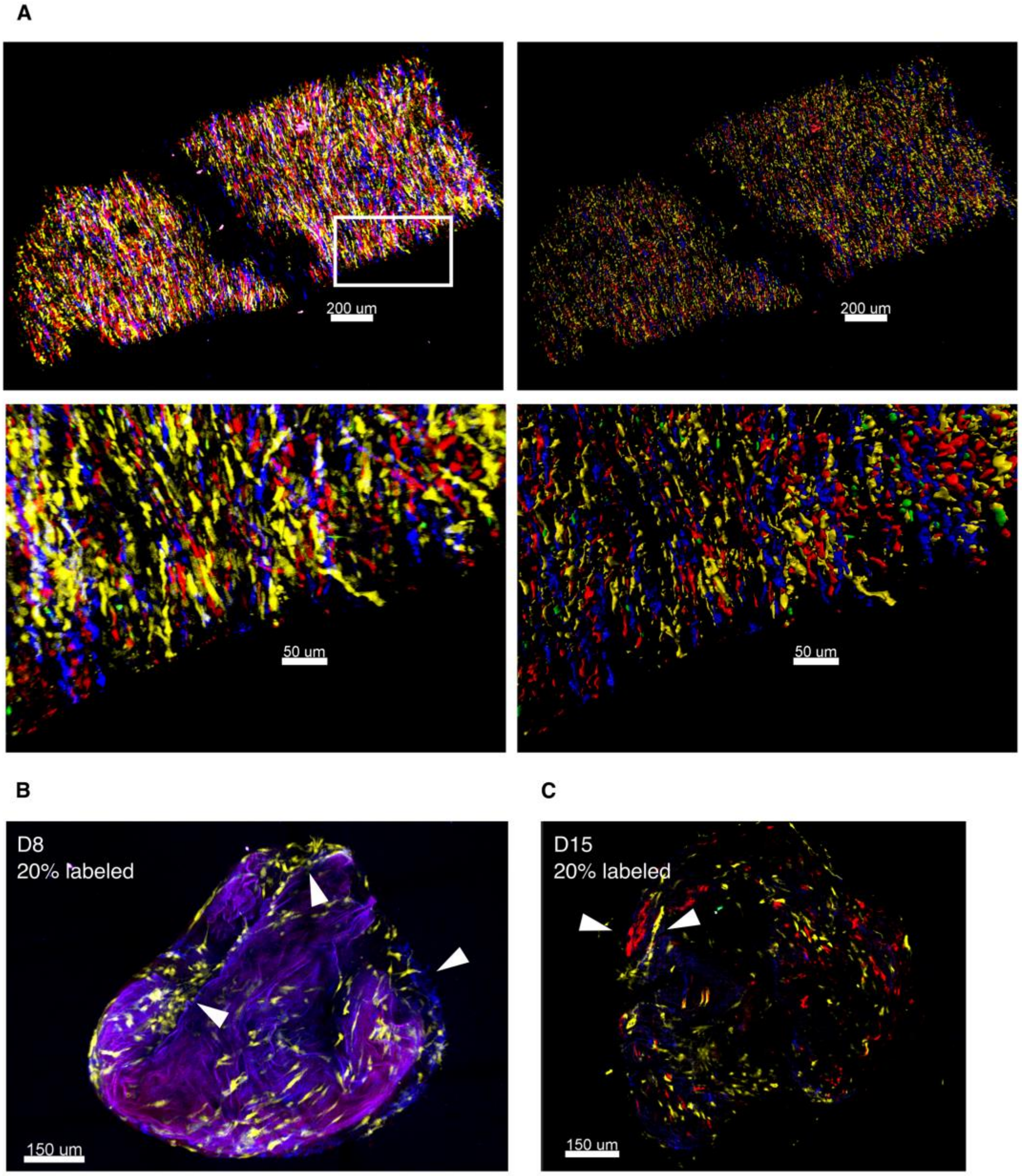


**Figure II: Tissue explant analysis. A,** Surface rendering in Imaris of confocal images for an aortic tissue explant analyzed at day 0. Signals for Confetti colors are shown and lower panels show magnified view of boxed region. Original images (left) and reconstructions after surface-rendering (right) are shown. Scalebar = 200 μm (top) or 50 μm (lower panels). **B, C**, Confocal images of tissue explants from Myh11-Confetti animals with lower frequency of VSMC-lineage labeling (20%, 2 injections of 0.1 mg tamoxifen) after culture for 8 (B, max projection) or 15 days (C, single Z-scan). Arrows point to mono-chromatic VSMC patches. Scalebar = 150 μm.

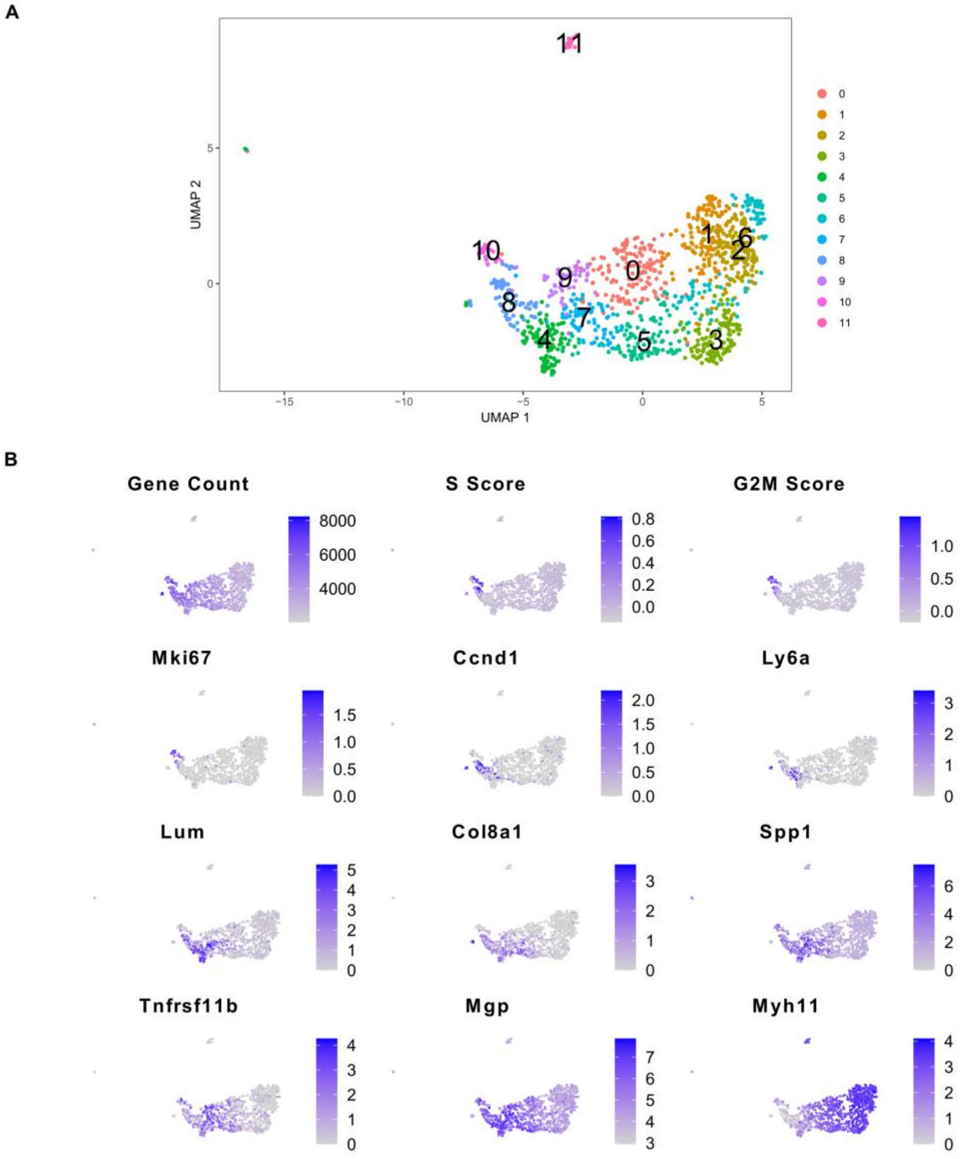


**Figure III: Feature plots of mouse D7 scRNA-seq dataset including cell cluster 11.** UMAP cluster **(A)** and feature plots **(B)** corresponding to Figure 4A, B including cluster 11. Cluster 11 markers include *Prss12*, *Prrx2*, *Rgs5* similar to the "minor SMC" population described by Pan et al (Pan et al., 2020).

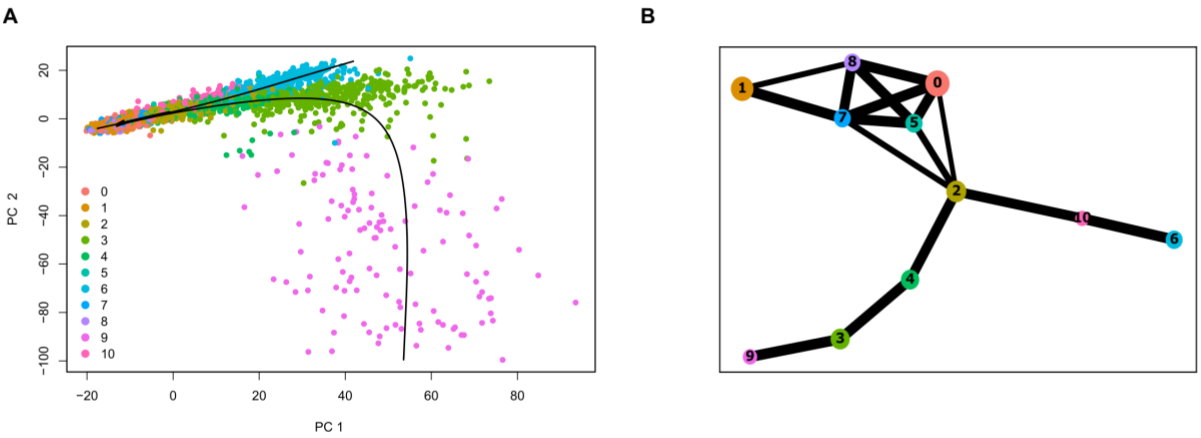


**Figure IV: Trajectory inference for mouse D5 scRNA-seq data.** Trajectory inference analyses of mouse D5 scRNA-seq dataset from VSMC-lineage label+ cells from Myh11-Confetti animals 5 days after carotid ligation (mouse D5 scRNA-seq dataset) using the slingshot package (A) or partition-based graph abstraction (PAGA, B). Cluster numbers and color-coding refer to clustering shown in Figure 5A of the main manuscript and are also indicated in panel A. **A,** Plot of Principal components 1 and 2 (PC1, PC2) where each dot represents a single cell. Black lines indicate slingshot identified paths; Path1: cluster 1 to cluster 9, and Path2: cluster 1 to cluster 6. **B,** PAGA connectivity map where black lines show connections between individual clusters and line thickness indicates the strength of the connection.

A

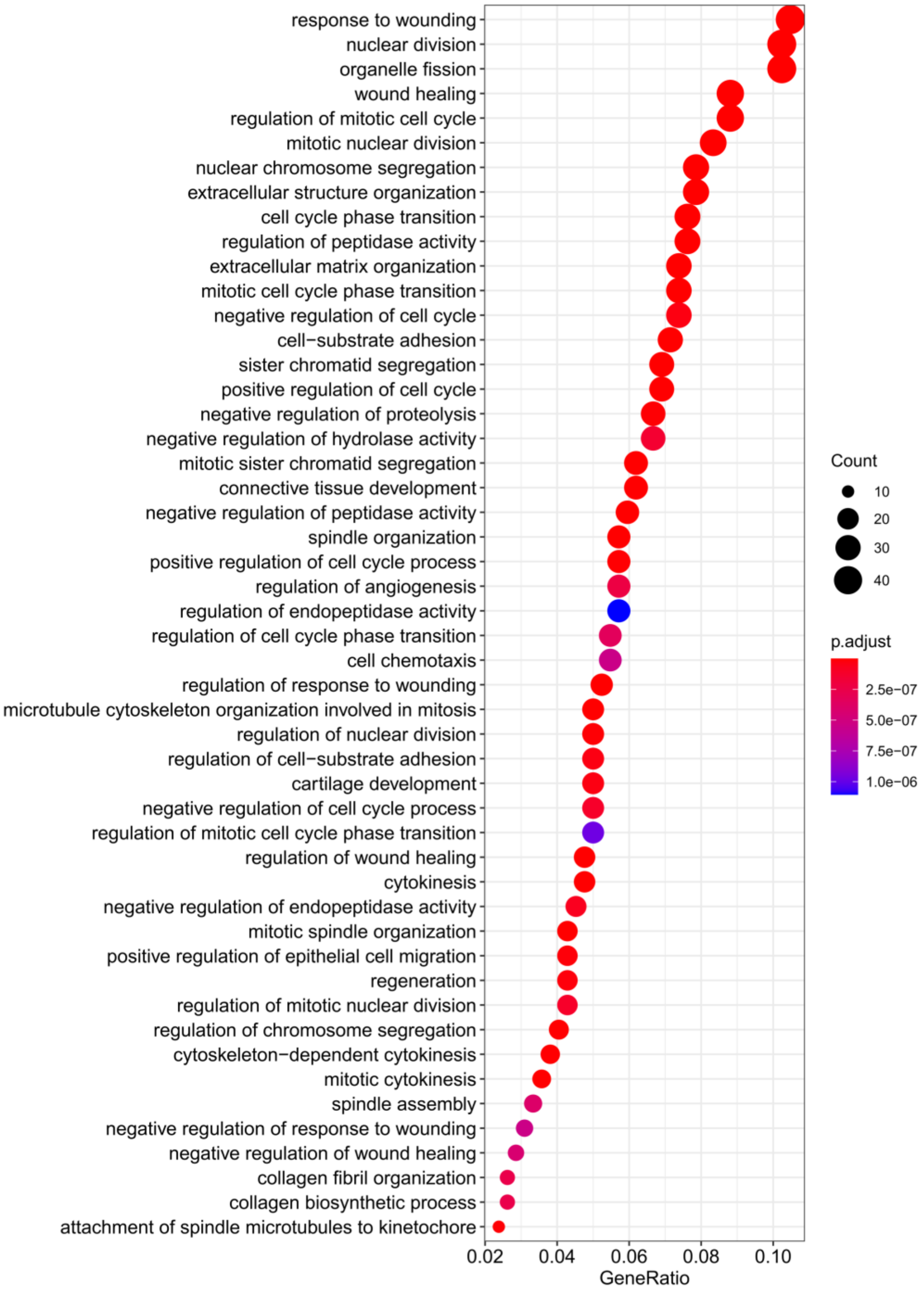
response to wounding
nuclear division
organelle fission
wound healing
regulation of mitotic cell cycle
mitotic nuclear division
nuclear chromosome segregation
extracellular structure organization
cell cycle phase transition
regulation of peptidase activity
extracellular matrix organization
mitotic cell cycle phase transition
negative regulation of cell cycle
cell−substrate adhesion
sister chromatid segregation
positive regulation of cell cycle
negative regulation of proteolysis
negative regulation of hydrolase activity
mitotic sister chromatid segregation
connective tissue development
negative regulation of peptidase activity
spindle organization
positive regulation of cell cycle process
regulation of angiogenesis
regulation of endopeptidase activity
regulation of cell cycle phase transition
cell chemotaxis
regulation of response to wounding
microtubule cytoskeleton organization involved in mitosis
regulation of nuclear division
regulation of cell−substrate adhesion
cartilage development
negative regulation of cell cycle process
regulation of mitotic cell cycle phase transition
regulation of wound healing
cytokinesis
negative regulation of endopeptidase activity
mitotic spindle organization
positive regulation of epithelial cell migration
regeneration
regulation of mitotic nuclear division
regulation of chromosome segregation
cytoskeleton−dependent cytokinesis
mitotic cytokinesis
spindle assembly
negative regulation of response to wounding
negative regulation of wound healing
collagen fibril organization
collagen biosynthetic process
attachment of spindle microtubules to kinetochore
0.02
0.04
0.06
0.08
0.10
GeneRatio
Count
10
20
30
40
p.adjust
2.5e−07
5.0e−07
7.5e−07
1.0e−06

**B**

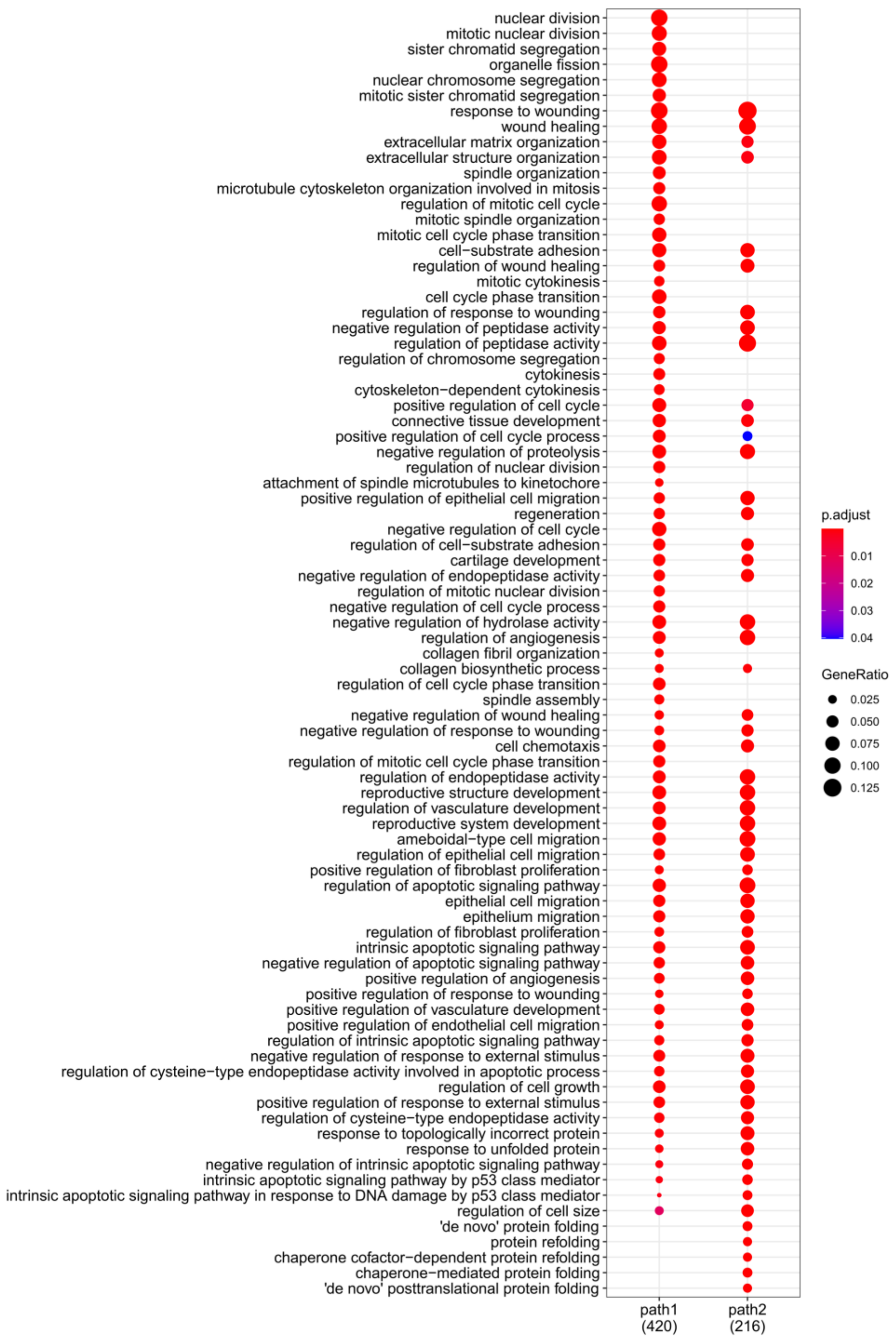


**Figure V: Gene ontology (GO) analysis of genes showing trajectory-induced expression in mouse D5 scRNA-seq dataset.** Selected enriched GO-terms for genes showing induced expression along path1 (gene clades 1, 2, 5, 7, 8, 10) **(A)** and induced expression along path1 (gene clades 1, 2, 5, 7, 8, 10) versus path2 (clade 2) **(B).**

A

B

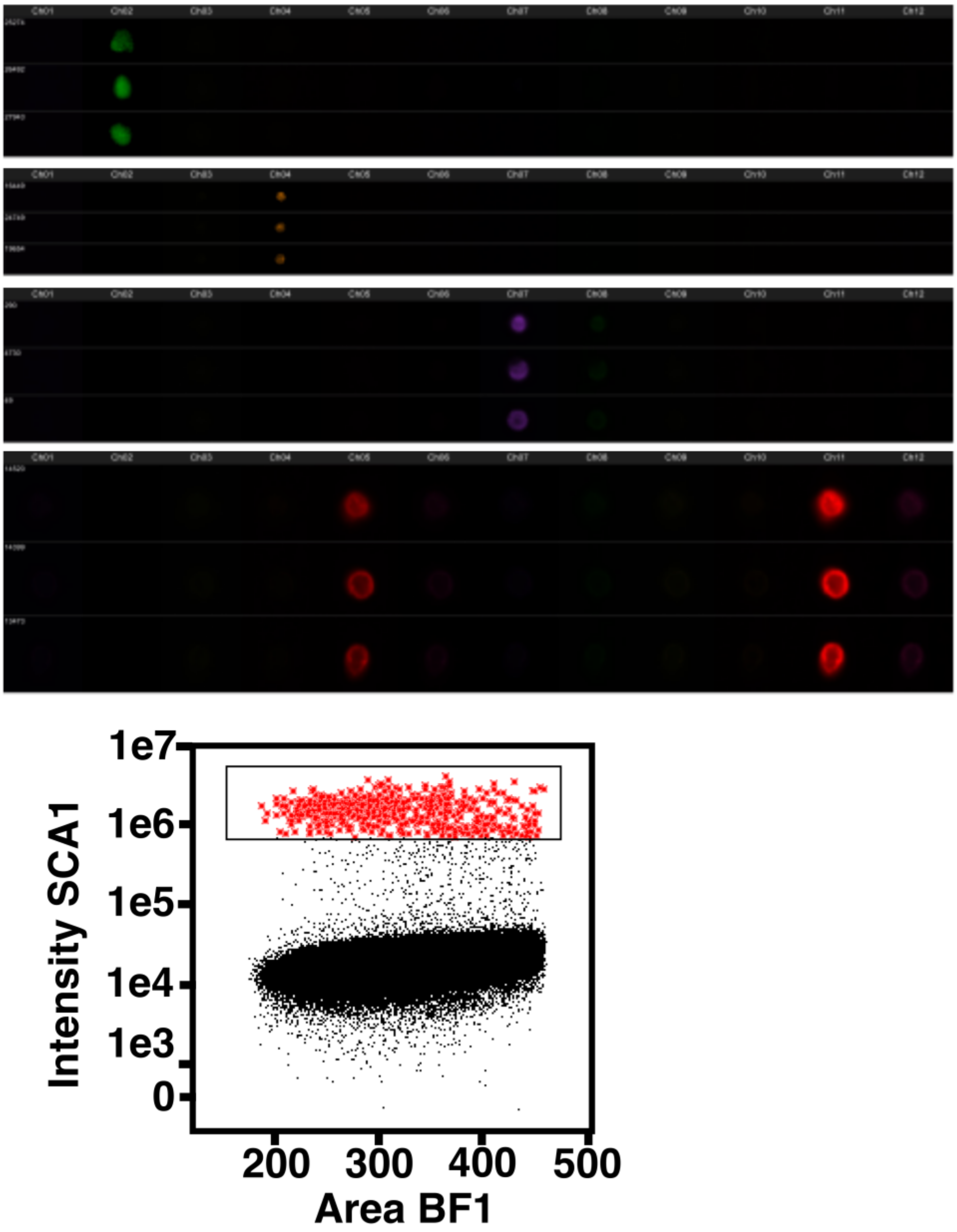


**Figure VI: ImageStream analysis. A,** Controls for ImageStream analysis showing unstained medial VSMCs (EYFP+, top) and adventitial cells (EYFP-) stained with anti-ROCK1 (orange), Phalloidin- iFluor™ 350 (magenta) or anti-SCA1 (red). SCA1 is detected in two channels. **B,** Dotplot showing intensity of SCA1 signal versus bright field area (BF1). Gate used to define SCA1+ cells in Figure 6 is indicated.

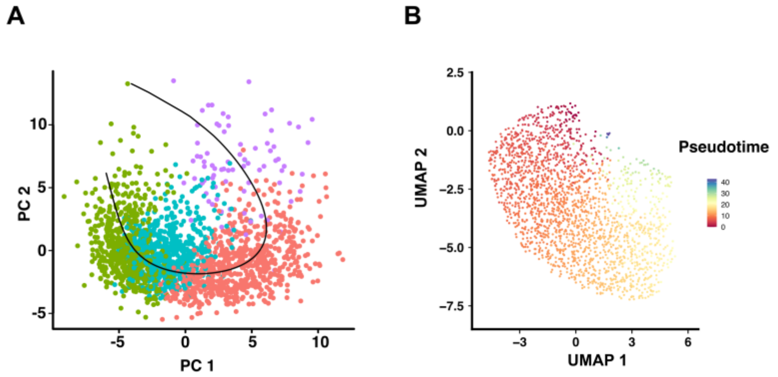


**Figure VII: Trajectory inference for human VSMC scRNA-seq dataset. A,** Principal component (PC) plot showing PC2 versus PC1 for human medial layer scRNA-seq dataset with cells color-coded by cluster identity (cluster 0 in red, cluster 1 in green, cluster 2 in turquoise and cluster 3 in purple). A black line shows the path from cluster 1 to 3 identified using trajectory analysis. **B,** UMAP of human medial layer scRNA-seq data with pseudotime for each cell shown using a scale from dark red to blue.

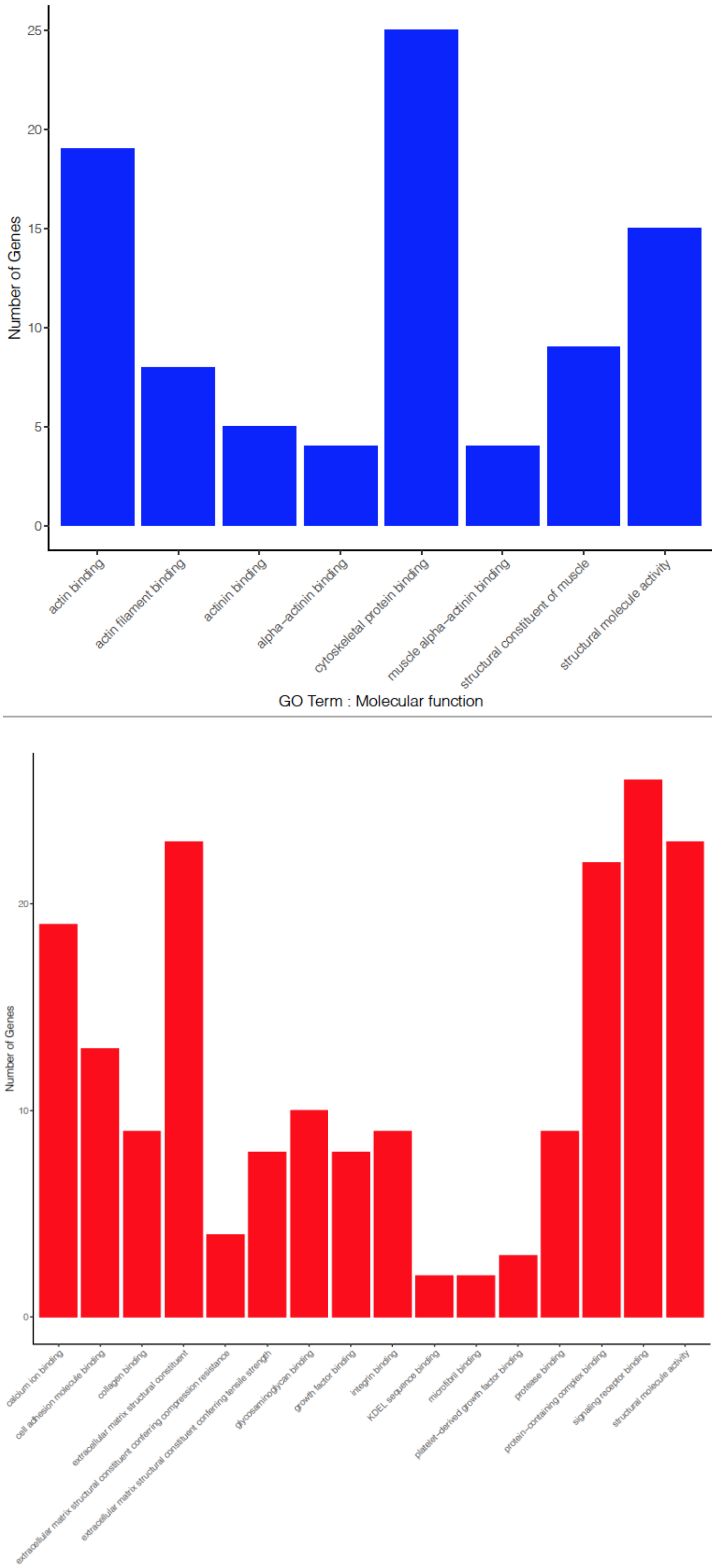


**Figure VIII: Enriched gene ontology terms for human pseudotime-associated genes.** Number of genes for enriched Molecular Function GO-terms in pseudotime-downregulated (top, blue) and pseudotime-induced genes (lower, red). Top 200 pseudotime-associated genes were used for this analysis.
